# Topological Learning for Brain Networks

**DOI:** 10.1101/2020.11.30.404665

**Authors:** Tananun Songdechakraiwut, Moo K. Chung

## Abstract

This paper proposes a novel topological learning framework that can integrate networks of different sizes and topology through persistent homology. This is possible through the introduction of a new topological loss function that enables such challenging task. The use of the proposed loss function bypasses the intrinsic computational bottleneck associated with matching networks. We validate the method in extensive statistical simulations with ground truth to assess the effectiveness of the topological loss in discriminating networks with different topology. The method is further applied to a twin brain imaging study in determining if the brain network is genetically heritable. The challenge is in overlaying the topologically different functional brain networks obtained from the resting-state functional magnetic resonance imaging (fMRI) onto the template structural brain network obtained through the diffusion tensor imaging (DTI).

## 1 Introduction

The human brain is an extraordinarily complicated system of interconnected neurons. The brain is roughly estimated to have 10^12^ neurons with 10^15^ synapses [1]. In an ideal situation, a brain network should be modeled using neurons as network nodes with every synapse as an edge connecting the nodes. Due to the technical limitation of magnetic resonance imaging (MRI) scanners, it is not possible to do so yet. In the popular brain network modeling framework, the whole brain is parcellated into *d* disjoint regions, where *d* is usually a few hundreds [2, 3, 4, 5, 6, 7, 8, 9, 10, 11, 12, 13, 14]. Subsequently, functional or structural information is overlaid on top of the parcellation to obtain *d* × *d* connectivity matrices that measure the strength of connectivity between brain regions for standard network analysis. For instance, the Automated Anatomical Labeling (AAL) partitions the brain into 116 regions (Figure 1) [2]. These disjoint brain regions form nodes in the brain network. Although there has been a long-standing effort toward standardizing the parcellation, there is no single universally accepted approach.

**Figure 1:**
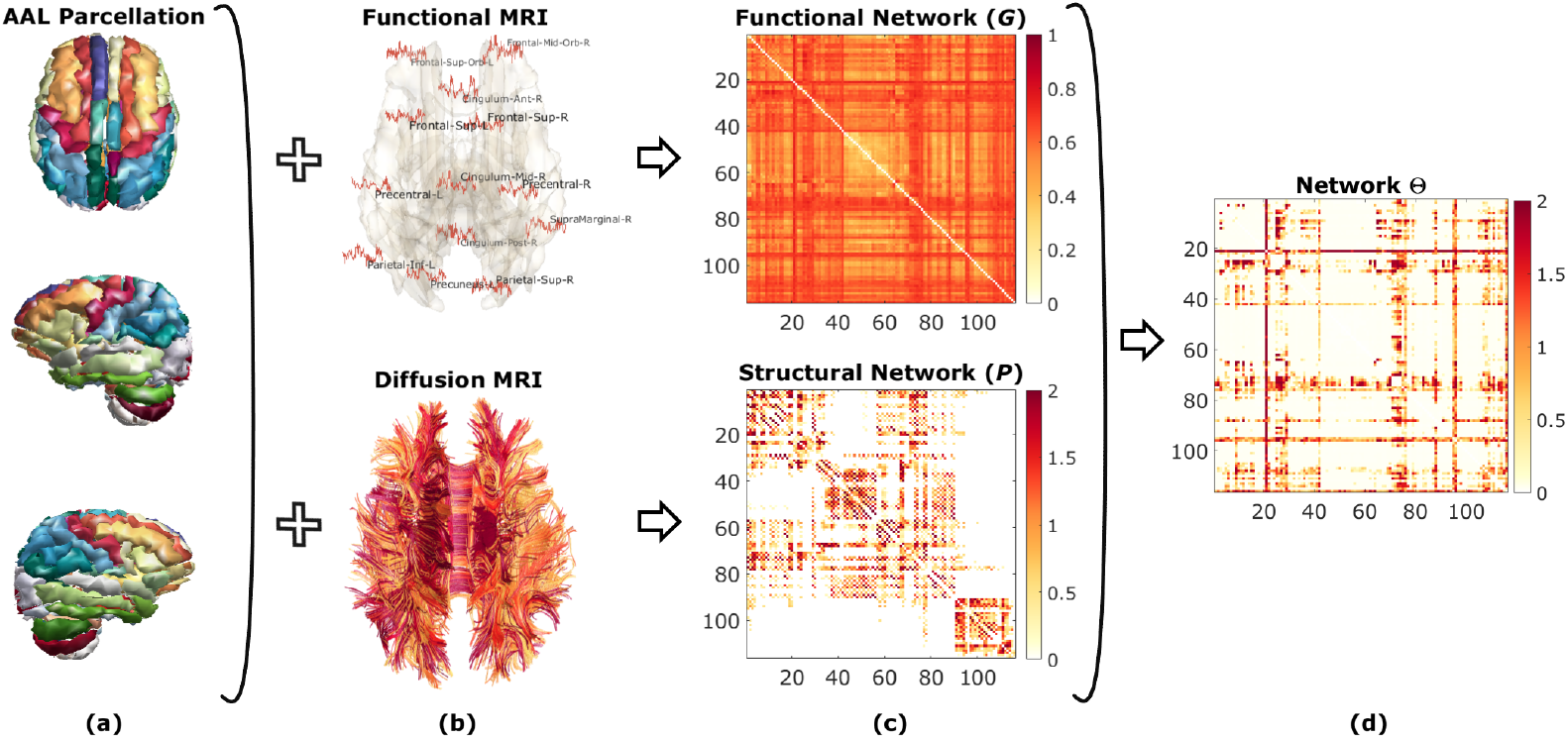
Schematic of topological learning on brain networks. (a) The Automated Anatomical Labeling (AAL) atlas obtained through structural-MRI is used to partition the human brain into 116 disjoint regions, which form the nodes of brain networks. In functional MRI, brain activity at each node is measured as a time series of changes associated with the relative blood oxygenation level (b-top). The functional connectivity between two nodes is given as the correlation between their fMRI time series, resulting in the functional network *G* through the metric transformation (c-top). The structural connectivity between two brain regions is measured by the number of white matter fiber tracts passing through them using diffusion MRI (b-bottom). Structural connectivities over all subjects are then normalized and scaled, resulting in the structural network *P* (c-bottom) that serves as the template where statistical analysis can be performed. The structural network *P* is sparse while the functional network *G* is densely connected and has more cycles. Since both networks are topologically different, it is difficult to integrate them together in a coherent model. Simply overlaying functional brain networks on top of the structural network, as usually done in the field, will completely destroy 1D topology (cycles) of the functional networks [15]. (d) Using the proposed framework, we learn network Θ that has the topological characteristics of both functional and structural networks.

Connectivity between brain regions that defines edges in the brain network is usually determined by the type of imaging modality [16]. Structural connectivity is obtained through diffusion MRI, which can trace the white matter fibers connecting brain regions. The strength of structural connectivity between the brain regions is determined by the number of fibers passing through them [17, 18]. The structural brain network is expected to exhibit sparse topology without many loops or cycles (Figure 1) [19, 17, 20]. On the other hand, functional connectivity obtained from the resting-state functional MRI (fMRI) is often computed as the Pearson correlation coefficient between brain regions [21, 22, 23, 24]. While structural connectivity provides information whether the brain regions are physically connected through white matter fibers, functional connectivity exhibits connections between two regions without the direct neuroanatomical connections through additional intermediate connections [25, 26]. Thus, resting-state functional brain networks are often very dense with thousands of cycles. Both structural and functional brain networks provide topologically different information. Existing graph theory based brain network analyses have shown that there is some common but limited topological profile that is conserved for both structural and functional brain networks [27]. However, due to the difficulty of integrating both networks in a coherent statistical framework, not much research has been done on integrating such networks at the localized edge level. Many multimodal network researchers focus on comparing summary graph theory features across different networks [27, 28, 29]. Although there are not many, some statistical studies have focused on fusing networks derived from both modalities *probabilistically*, which can easily destroy the aforementioned topological difference of the networks [30, 31]. Thus, there is a need for a new multimodal network model that can easily integrate networks of different topology at the localized connection level.

Topological Data Analysis (TDA) [32, 33], a general framework based on algebraic topology, can provide such novel solution to the long-standing multimodal brain network analysis challenge. Numerous TDA studies have been applied to increasingly diverse problems such as genetics [34, 24], epileptic seizure detection [35], sexual dimorphism in the human brain [36], analysis of brain arteries [37], image segmentation [38], classification [39, 40, 41], clinical predictive model [42] and persistence-based clustering [43]. Persistent homology begins to emerge as a powerful mathematical representation to understand, characterize and quantify topology of brain networks [44, 24]. In persistent homology, topological basis such as connected components and cycles are measured across different spatial resolutions. As the resolution changes, such features are born and die. Persistent homology associates the life-time to these features in the form of 1D interval from birth to death. Long-lived features persist over a long range of resolutions and are considered as signal while short-lived features are considered as noise [45]. The collection of such intervals is summarized as a *barcode*, which completely characterizes the topology of underlying data [46].

Recently, it was proposed to penalizes barcodes through *topological loss* for image segmentation [47, 48]. While the approach allows to incorporate topological information into segmentation problem, the method has been limited to 2D image segmentation with a small number of topological features due to its brute-force computation involving *O*(*d*^6^) run time [49, 50]. Barcodes are typically computed at a finite set of pre-specified resolutions. A sufficient number of such resolutions is required to give a reasonably accurate estimation of barcodes, which quickly increases computational complexity when the size of data increases [51, 48]. This is impractical in brain networks with far larger number of topological features involving hundreds of connected components and thousands of cycles. In this paper, motivated by [47, 48], we propose a more principled approach that learns the topological structure of brain networks with large number of features in *O*(*d*^2^ log *d*) run time. Our proposed method bypasses the intrinsic computational bottleneck and thus enables us to perform various topology computations and optimizations at every possible resolution.

We illustrate the method in studying the resting-state functional MRI networks of 194 twin pairs obtained from the Human Connectome Project [52, 53]. HCP twin brain imaging data is considered as the *gold standard*, where the zigosity is confirmed by the blood and saliva test [54]. Monozygotic (MZ) twins share 100% of genes while dizygotic (DZ) twins share 50% of genes [55, 56]. MZ-twins are more similar or concordant than DZ-twins for cognitive aging, cognitive dysfunction and Alzheimer’s disease [57]. These genetic differences allow us to pull apart and examine genetic and environmental influences easily *in vivo*. The difference between MZ- and DZ-twins directly quantify the extent to which phenotypes are influenced by genetic factors. If MZ-twins show more similarity on a given trait compared to DZ-twins, this provides evidence that genes significantly influence that trait.

Previous twin brain imaging studies mainly used univariate imaging phenotypes such as cortical surface thickness [58], fractional anisotropy [59], functional activation [60, 61, 62] in determining heritability in few regions of interest. Compared to existing studies on univariate imaging phenotypes, there are not many studies on the heritability of the whole brain functional networks [60]. Measures of network topology and features may be worth investigating as intermediate phenotypes that indicate the genetic risk for a neuropsychiatric disorder [63]. However, the brain network analysis has not yet been adapted for this purpose beyond a small number of regions. Determining the extent of heritability of the whole brain networks is the first necessary prerequisite for identifying network-based endophenotypes. Using the proposed method, we propose to determine the heritability of functional brain networks while integrating the structural brain network information. We demonstrate that our method increases the sensitivity in detecting subtle genetic signals.

## 2 Methods

### Graph filtration

We consider a network, which is represented as a weighted graph *G* = (*V, w*) comprising a set of nodes *V* and unique positive symmetric edge weights *w* = (*w_ij_*). We assume the graph has no self-loops or parallel edges. The cardinality of sets is denoted using |·|. The number of nodes and edges are then denoted as |*V*| and |*E*|. Since *G* is a weighted complete graph, we have |*E*| = |*V*|(|*V*| − 1)/2. The binary graph *G_ϵ_* = (*V*, *w_ϵ_*) of *G* is a graph consisting of node set *V* and binary edge weight *w_ϵ_* given by

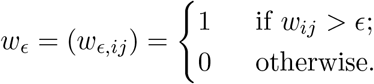

*G*_0_ is a complete graph while *G*_∞_ is the node set *V*. The binary network *G_ϵ_* is the 1-skeleton, which is a simplicial complex consisting of nodes and edges only. In 1-skeleton, 0-dimensional (0D) holes are connected components and 1-dimensional (1D) holes are cycles [24]. Unlike Rips complex, which is the usual basic building block of persistent homology, there are no more higher dimensional topological features to compute beyond 1D in 1-skeleton. The number of connected components and the number of *independent* cycles in the network are referred to as the 0-th Betti number and the 1-st Betti number, respectively. The *i*-th Betti number of network *G_ϵ_* is denoted as *β_i_*(*G_ϵ_*).

A graph filtration of *G* is defined as a collection of nested binary networks [44, 64, 34, 24]:

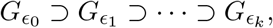

where *ϵ*_0_ < *ϵ*_1_ <… < *ϵ_k_* are filtration values. Since there are many possible graph filtrations over the different choice of filtration values, we will simply take edge weights as filtration values and make the graph filtration unique.

By increasing *ϵ*, we are thresholding at higher connectivity, resulting in more edges being removed. Each removed edge either increments the number of connected components (*β*_0_) or decrements the number of cycles (*β*_1_) *at most by one* [24]. Figure 2-a displays an example of the graph filtration on a four-node network. Note *G*_0_ is a complete graph consisting of a single connected component, while *G*_∞_ is a node set. Thus, *β*_0_ is monotonically increasing from *β*_0_(*G*_0_) = 1 to *β*_0_(*G*_∞_) = |*V*| and the increment is at most by one.

**Figure 2:**
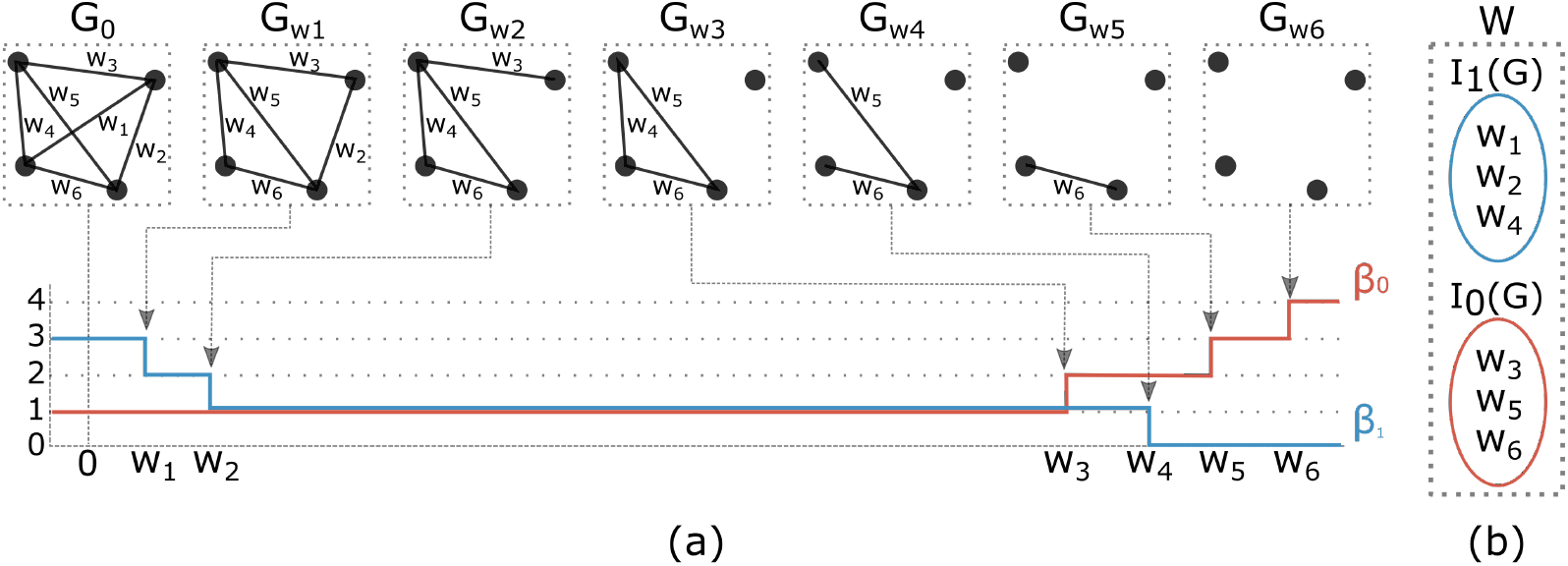
(a) Graph filtration on four-node network *G*. *β*_0_ is monotonically increasing while *β*_1_ is monotonically decreasing over the filtration. Connected components are born at the edge weights *w*_3_, *w*_5_, *w*_6_ while cycles die at the edge weights *w*_1_, *w*_2_, *w*_4_. A cycle consisting of *w*_4_, *w*_5_, *w*_6_ persists longer than any other 1D features and considered as topological signal. 0D barcode *P*_0_ = {(−∞, ∞), (*w*_3_, ∞), (*w*_5_, ∞), (*w*_6_, ∞)} is represented using the birth values as *I*_0_(*G*) = {*w*_3_, *w*_5_, *w*_6_}. 1D barcode *P*_1_ = {(−∞, *w*_1_), (−∞, *w*_2_), (−∞, *w*_4_)} is represented using the death values as *I*_1_(*G*) = {*w*_1_, *w*_2_, *w*_4_}. (b) We can prove that 0D and 1D barcodes uniquely partition the edge weight set *W*, i.e., *W* = *I*_0_(*G*) ∪ *I*_1_(*G*) with *I*_0_(*G*) ∩ *l*_1_(*G*) = ∅.

Similarly, *β*_1_ is monotonically decreasing from *β*_1_(*G*_0_) to *β*_1_(*G_∞_*) and the decrement is at most by one. *β*_1_(*G_∞_*) is trivially zero. How many cycles are there in the complete graph *G*_0_? The Euler characteristic χ of network *G*_0_ is given by [24]

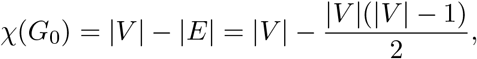

which can also be written as *χ*(*G*_0_) = *β*_0_(*G*_0_) − *β*_1_(*G*_0_) [65]. Thus, there are

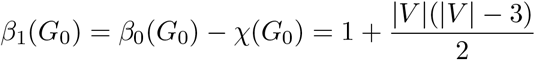

number of cycles. In Figure 2-a, *G*_0_ is a tetrahedron with 4 faces, which are cycles. However, there exists only 3 independent cycles and *β*_1_(*G*_0_) = 3.

### Barcodes in graph filtration

Persistent homology keeps track of appearances (birth) and disappearances (death) of connected components and cycles over the whole range of filtration values *ϵ*, and associates their *persistence* (the life-time measured as the duration of birth to death) to them. Longer persistence indicates the presence of topological signal while shorter persistence is interpreted as topological noise [66]. One representation to capture the persistence of such topological features is in the form of barcodes [46, 65, 67]. The 0D and 1D barcodes corresponding to the network *G* are collections *P*_0_ and *P*_1_ of intervals [*ϵ_b_*, *ϵ_d_*] such that each interval tabulates the life-time of a connected component and a cycle, respectively, that appears at the first filtration value *ϵ_b_* and vanishes at the last filtration value *ϵ_d_* (Figure 2-a).

Since *G*_0_ is a complete graph, a single connected component, we simply treat *G*_0_ to be born at −∞. By the monotonicity of *β*_0_ [24], the number of connected components is non-decreasing as *ϵ* increases. Once a component is born, it never dies so we have death value *ϵ_d_* = ∞ for every connected component. Ignoring one birth value corresponding to the complete graph at −∞, the total number of birth values of connected components is then

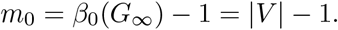

Thus, we simply represent 0D barcode of the network only using the set of increasing birth values

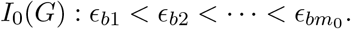

Since *G*_0_ is the complete graph, all cycles are considered to be born at *ϵ_b_* = −∞. By the monotonicity of *β*_1_, the number of cycles in 1D barcode is nonincreasing as *ϵ* increases. Thus, a cycle will die one at a time over increasing filtration values. The total number of death values of cycles is then equal to the number of cycles:

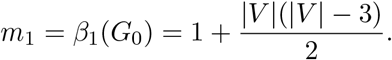

Similarly, we can also represent 1D barcode only using the set of increasing death values

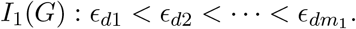

Figure 2-a illustrates how 0D and 1D barcodes corresponding to a four-node network are simplified to a collection of birth and death values respectively.

Under the graph filtration, deleting edge *w_ij_* in the network *G* must result in either the birth of a connected component or the death of a cycle. The birth of a component and the death of a cycle cannot possibly happen at the same time. Every edge weight must be in either 0D barcode or 1D barcode (Figure 2-b). Thus, we have

#### Theorem 1.

*The set of 0D birth values I*_0_(*G*) *and 1D death values I*_1_(*G*) *partition the edge weight set W such that W* = *I*_0_(*G*) ∪ *I*_1_(*G*) *with I*_0_(*G*) ∩ *l*_1_(*G*) = ∅.

Theorem 1 is a non-trivial statement and used in the development of the proposed topological learning framework.

### Topological loss

Since a network is topologically completely characterized by 0D and 1D barcodes, the topological similarity between two networks can be measured using differences of such barcodes. We modified the Wasserstein distance to measure the differences in the barcodes for 1-skeleton [47, 48, 38]. Let Θ = (*V*^Θ^, *w*^Θ^) and *P* = (*V^P^*, *w^P^*) be two networks. For now, we will simply assume that the two networks have the same size, i.e., |*V*^Θ^| = |*V^P^*| but the method works for |*V*^Θ^| ≠|*V^P^*| as well through the data augmentation (explained later). The topological loss 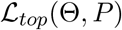 is defined as the optimal matching cost

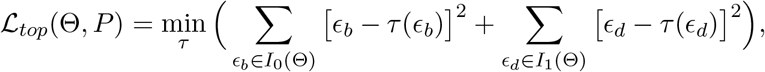

where *τ* is a *bijection* from *I*_0_(Θ) ∪ *I*_1_(Θ) to *I*_0_(*P*) ∪ *I*_1_(*P*). It is reasonable to match 0D to 0D persistences and 1D to 1D persistences. Thus, we further restrict the bijection to map from *I*_0_(Θ) to *I*_0_(*P*) and *I*_1_(Θ) to *I*_1_(*P*). Subsequently, it is equivalent to optimize separately as

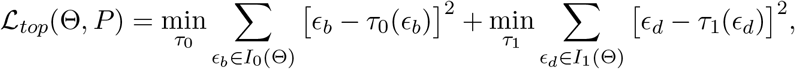

where *τ*_0_ is a bijection from *I*_0_(Θ) to *I*_0_(*P*) and *τ*_1_ is a bijection from *I*_1_(Θ) to *I*_1_(*P*). We will call the first term as 0D topological loss, which measures the topological similarity between the two networks Θ and *P* using the difference in 0D barcodes and is denoted by 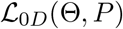. Similarly, we will call the second term as 1D topological loss, which measures the topological similarity through the difference of 1D barcodes and is denoted by 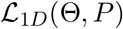. The 0D topological loss 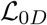 is a variation to the standard *assignment problem* and usually solved in a greedy fashion using Hungarian algorithm in *O*(|*I*_0_(Θ)|^3^), or equivalently *O*(|*V*^Θ^|^3^), in combinatorial optimization [49]. However, for 1-skeletons, minimum matching is given exactly and can be numerically computed in *O*|*I*_0_(Θ)| log |*I*_0_(Θ)|) time:

#### Theorem 2.

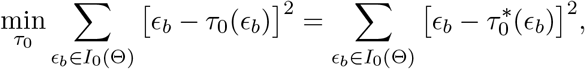

*where* 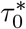 *maps the i-th smallest birth value in I*_0_(Θ) *to the i-th smallest birth value in I*_0_(*P*) *for all i.*

The minimization in Theorem 2 is equivalent to the following assignment problem. For monotonic sequences

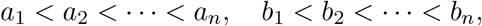

we consider finding 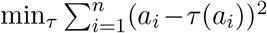 over all possible bijection *τ*. The minimization is equivalent to maximizing the assignment cost 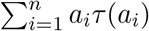. Then, we follow an inductive argument. For *n* = 2, there are two possible assignments: *a*_1_*b*_1_ + *a*_2_*b*_2_ or *a*_1_*b*_2_ + *a*_2_*b*_1_. Since *a*_1_*b*_1_ + *a*_2_*b*_2_ *> a*_1_*b*_2_ + *a*_2_*b*_1_, *τ* (*a_i_*) = *b_i_* is the optimal matching. For *n* = *k*, we assume *τ* (*a_i_*) = *b_i_* is the optimal matching for all *i*. For *n* = *k* + 1, the cost is given by

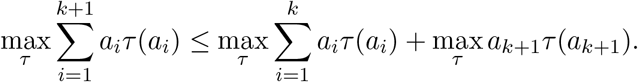

The first term is maximized if *τ*(*a_i_*) = *b_i_*. The second term is maximized if *τ*(*a*_*k*+1_) = *b*_*k*+1_. Thus, we have proved the statement. The immediate consequence of the theorem is that the loss function 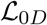 is symmetric: 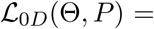 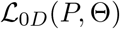.

Similarly for the 1D topological loss 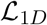, we also have

#### Theorem 3.

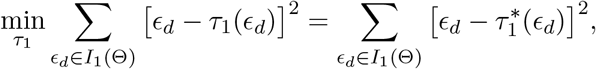

where 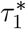 *maps the i-th smallest death value in I*_1_(Θ) *to the i-th smallest death value in I*_1_(*P*) *for all i.*

Subsequently, the loss function 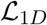 is symmetric: 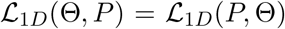 Similar to 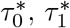 can be computed in *O |I*_1_(Θ)| log |*I*_1_(Θ)| time.

### Augmentation: networks with different sizes

When networks have different sizes in terms of the number of nodes, we can still find the optimal matching through the data augmentation on smaller networks. Similar data augmentations are done in matching various topological features of different sizes [48], matching trees of different sizes [68] or matching point sets of different sizes [24].

Let Θ = (*V* ^Θ^, *w*^Θ^) and *P* = (*V ^P^, w^P^*) be two networks of different sizes, i.e., |*V* ^Θ^|*/*= |*V ^P^*|. We have two cases as follows.

*Case* |*V*^Θ^| > |*V^P^*|: Since |*I*_0_(Θ)| > |*I*_0_(*P*)|, some birth values in Θ may not have corresponding matches in *P*. We match the first |*I*_0_(*P*)| smallest birth values in Θ to all the birth values in *P* in ascending order. Then, we match the remaining unmatched, large birth values in Θ to the largest edge weight in *P*. Connected components that are born later (larger birth values) have shorter persistence (life-times), which are considered as topological noise [46, 38, 48]. Thus, we need to map the noisy, unmatched connected components in Θ to the largest edge weight in *P* and identify 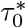. Similarly, we match the first *I*_1_(*P*) largest death values in Θ to all the death values in *P* and identify 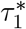.

*Case* |*V*^Θ^| < |*V^P^*|: Since |*I*_0_(Θ)| < |*I*_0_(*P*)|, some birth and death values in *P* may not have corresponding matches in Θ. Similar to the case |*V*^Θ^| > |*V*^*P*^, any unmatched birth or death values in *P* are matched to the largest or smallest edge weights in Θ respectively.

### Topological Learning

Most learning tasks such as regression, classification and clustering can be performed by minimizing the topological loss. Let *G*_1_ = (*V, w*^1^), *, G_n_* = (*V, w^n^*) be the observed networks with identical node set *V* that will be used as *training networks*. Let *P* = (*V ^P^, w^P^*) be a network expressing a prior topological knowledge. In brain network analysis [69, 15, 31], *G_k_* can be the functional brain network of the *k*-th subject obtained from the resting-state fMRI and *P* can be the template structural brain network obtained through DTI, where functional brain networks are often overlaid (Figure 1). The node sets *V* and *V ^P^* may differ. This can happen if we try to integrate brain networks obtained from different parcellations and studies.

We are interested in learning the model Θ = (*V*^Θ^, *w*^Θ^) from the set of training data. At the subject level, we train the model using an individual network *G_k_* by minimizing

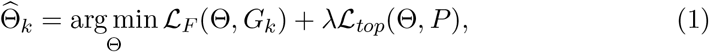

where the squared Frobenius loss 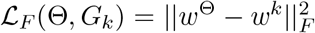 is the goodness-of-fit term between the model and the individual observation. The parameter *λ* controls the amount of topological information of network *P* we are introducing to the model. The larger the value of *λ*, the more we are learning toward *P*. If *λ* = 0, we no longer learn the topology of *P* but simply fit data to the individual network *G_k_*. The total loss is the sum of quadratic losses and thus it obtains the global minimum. Figure 3 shows the total loss function with the global minimum at *λ* = 0.9997 for a subject. The average global minimum across all subjects is obtained at *λ* = 1.0000±0.0002 showing highly stable result. Thus, we simply use *λ* = 1.0000 for all subjects. Such stable result is not possible in non-topological loss functions. Learned individual function networks are used in determining the genetic heritability in the Application section.

**Figure 3:**
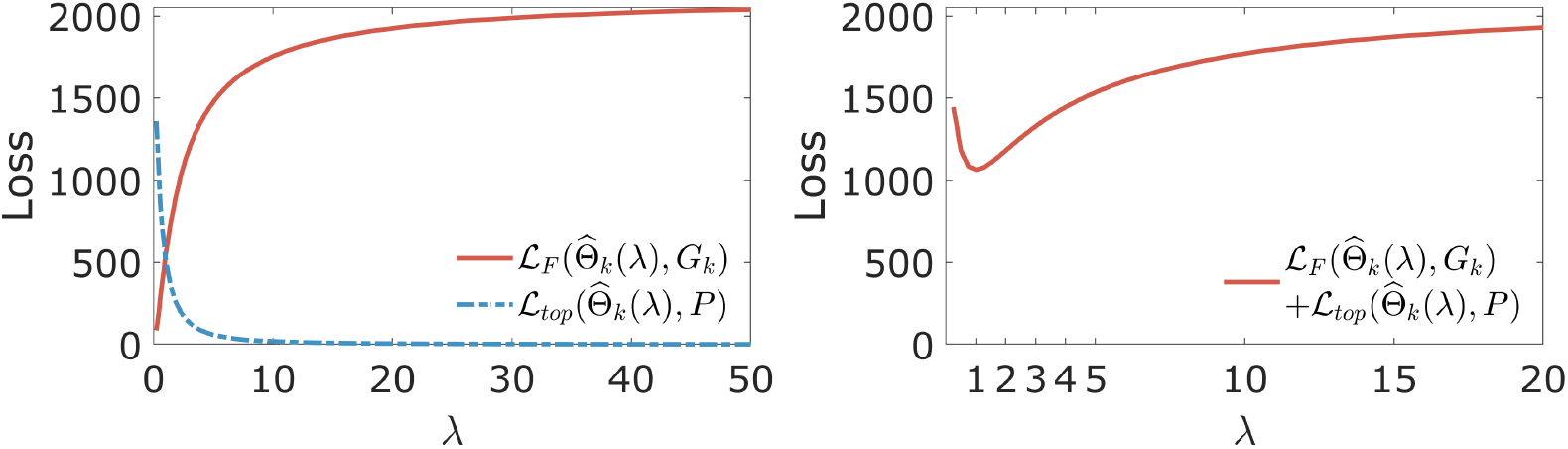
Topological learning on a functional brain network of a subject. When *λ* = 0, 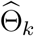 is simply equal to *G_k_*. As *λ* increases, 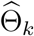 is deformed such that the topology of 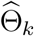 is closer to *P*. For this subject, the total loss function obtains the global minimum at *λ* = 0.9997. Other subjects show almost identical patterns.

Although the group level network learning is not the focus of this study, we can also learn network 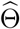 using all the training data such that

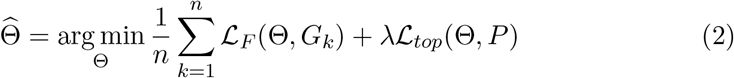

Figure 4 displays the average networks of females and males by minimizing the objective function (2) with different *λ* = 0, 1 and 100. Larger the value of *λ*, we are reinforcing the topology of structural brain network that is sparse onto the functional brain network (Figure 5). Even though we will not show in this paper, the statistical significance of network differences between females and males can be determined by using the exact topological inference developed in our previous work [56, 70].

**Figure 4:**
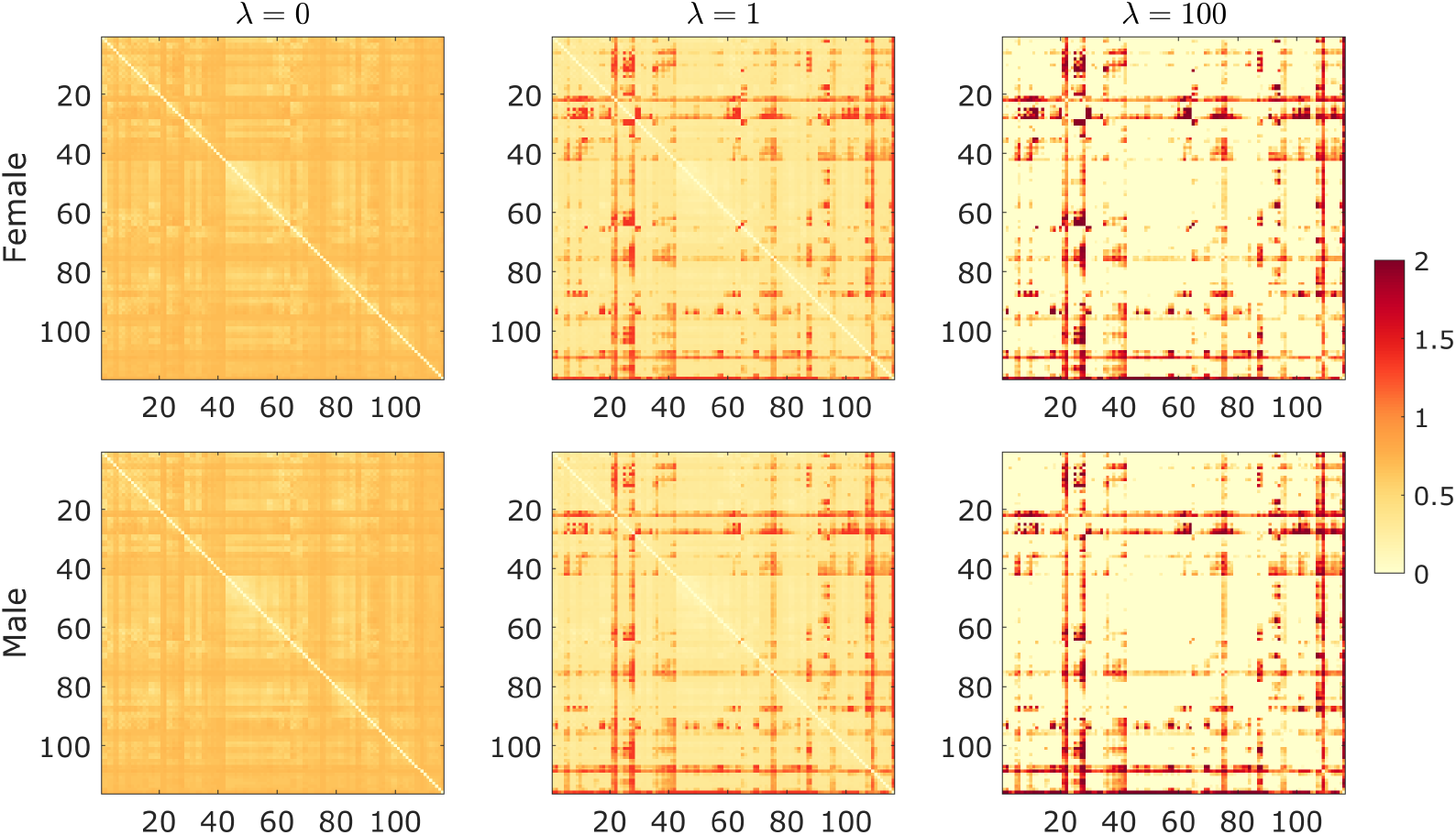
Group level networks of female (top row) and male (bottom row) are estimated by minimizing the objective function (2) with different *λ* = 0, 1 and 100.

**Figure 5:**
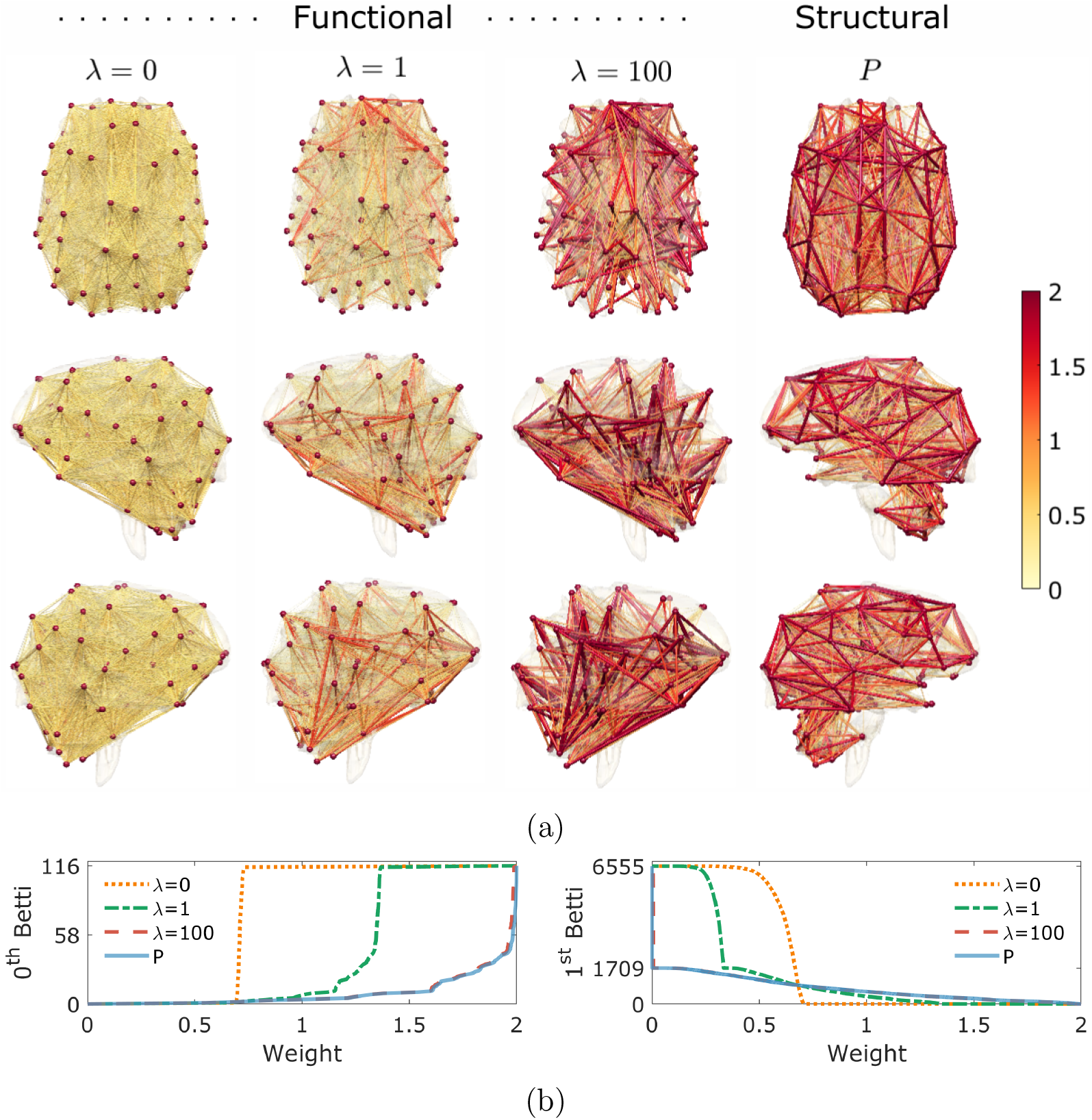
(a) The group level networks learned by minimizing the objective function (2) over all subjects with different *λ* = 0, 1 and 100. The template structural network *P* is shown in the last column. (b) As *λ* increases, Betti-plots of the group level network are adjusted toward that of *P*. *β*_0_-plot shows that the connected components in the structural network *P* are gradually born over wide range of edge weights during graph filtration. *β*_1_-plot shows topological sparsity of lack of cycles in the structural network *P*. While the group level network (when *λ* = 0) is densely connected with maximum 6555 number of cycles, the structural network is sparsely connected with only 1709 cycles.

### Averaging networks of different sizes and topology

As an application of the proposed topological learning framework, it is also possible to average networks of different sizes and topology directly without aligning to the template network as in (2). This might be useful in a situation when we do not have the template or it is not necessary to align networks to a template. Averaging networks of different topology is a difficult task using existing methods.

Given *n* networks *G*_1_ = (*V*_1_, *w*^1^), *, G_n_* = (*V_n_, w^n^*) with different node sets, we are interested in obtaining its average, which we will call the *topological mean*. Since the size and topology of the networks are different, we cannot simply average edge weight matrices *w*^1^, … *, w^n^* directly. Motivated by Fréchet mean [71, 72, 73], we obtain the topological mean 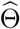 by minimizing the sum of topo-logical losses

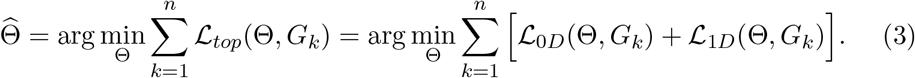

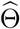 is viewed as a network that is the topological center of *n* networks. The optimization can be done analytically as follows.

For now, we assume the same number of nodes in the networks, which gives the same number *m*_0_ of birth values. The 0D topological loss 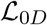 depends on the birth values of *G*_1_*,…, G_n_*. Let 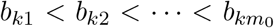be the birth values of network *G_k_*. Let 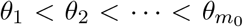 be the birth values of network Θ. By Theorem 2, the first term is equivalent to

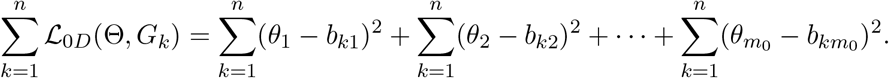

This is a quadratic so we can simply find the minimum by setting its derivative equal to zero. This is given by 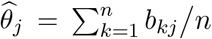. For the second term, it is similarly represented as the sum of squared differences of death values (Theorem 3). Thus, the birth and death values of the topological mean network Θ are simply given by the mean of all the birth and death values of *n* networks. Given all the birth and death values of a network, we can completely recover the topology of the network. For networks with different number of nodes, we augment birth and death values. Figure 6 illustrates toy examples of averaging networks of different sizes and topology.

**Figure 6:**
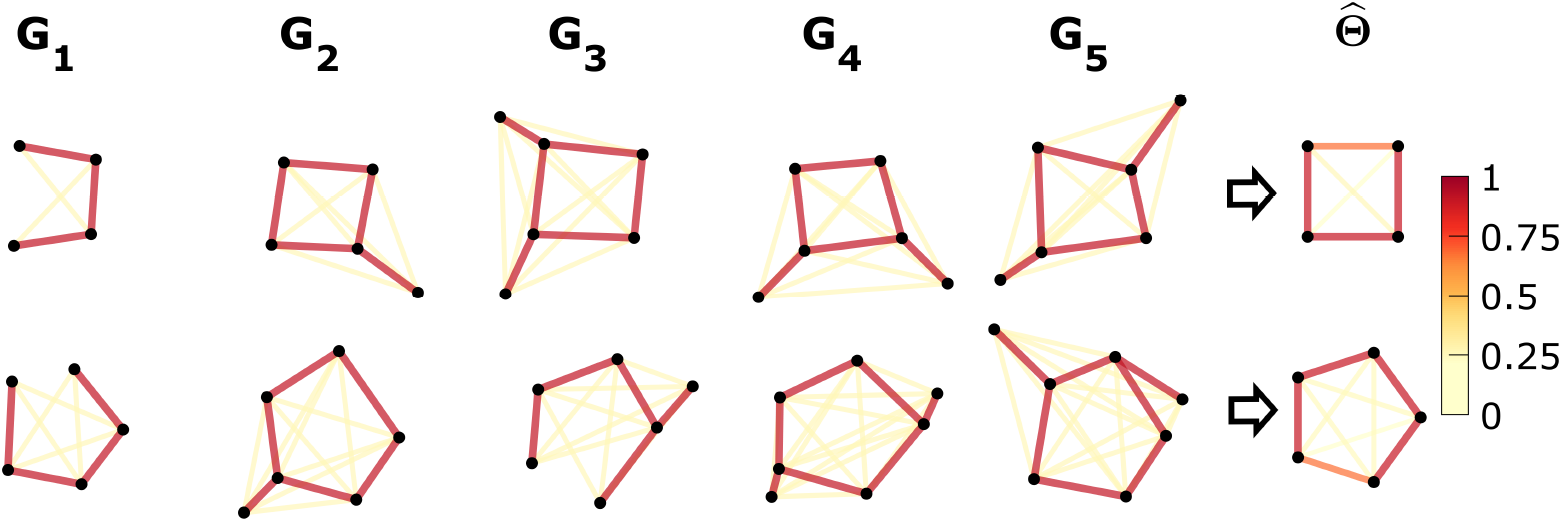
Examples of averaging networks of different sizes and topology using the proposed topological averaging. The topological mean network 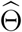 (right) is the topological center of five networks *G*_1_, *…, G*_5_ (left), showing the average topological pattern. The topological mean network 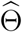 estimated by minimizing the sum of topological losses 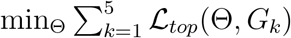. The topological mean network 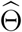 highlights topological characterization of the five networks. The existing methods will have difficulty averaging networks of different sizes and topology.

### Numerical implementation: gradient descent

The topological learning (1) estimates Θ = (*V* ^Θ^, *w*^Θ^) iteratively through gradient descent [74]. The gradient of the topological loss can be computed efficiently without taking numerical derivatives and its computation mainly comprises the computation of barcodes *I*_0_ and *I*_1_, and finding the optimal matchings through Theorems 2 and 3. The gradient of the topological loss 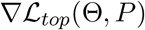 with respect to edge weight 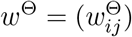 is given as a gradient matrix whose *ij*-th entry is

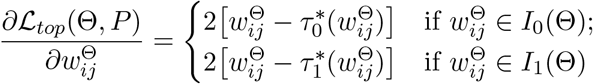

since *I*_0_(Θ) and *I*_1_(Θ) partition the weight set (Theorem 1). Intuitively, by slightly adjusting the edge weight 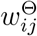, we have the slight adjustment of either a birth value in 0D barcode or a death value in 1D barcode, which slightly changes the topology of the network. During the estimation of Θ, we take steps in the direction of negative gradient such that

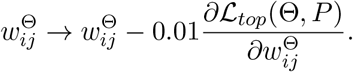

As 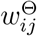 are moved closer to its optimal match, the topology of the estimated network Θ gets closer to that of *P* while the Frobenius norm keeps the estimation Θ close to the observed networks *G_k_*.

Finding 0D birth values *I*_0_ is *equivalent* to finding the sorted edge weights of the *maximum spanning tree* (MST) [44]. Once *I*_0_ is computed, *I*_1_ is simply given as the rest of the remaining edge weights (Theorem 1). Then, we can compute the optimal matchings 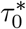 and 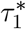 between Θ and *P* by simply sorting edge weights in the ascending order and matching them. The computational complexity for the topological loss gradient is dominated by the computation of the MST using the popular algorithms such as Prim’s and Kruskal’s. This takes *O*(|*E*^Θ^| log |*V* ^Θ^|) if |*V*^Θ^| ≥ |*V^P^*| or *O*(|*E^P^*| log |*V ^P^*|) if |*V*^Θ^| < |*V^P^*|.

## 3 Validation

For validation, we performed two simulation studies to assess the performance of the topological loss as a similarity measure between networks of different topology. Networks of the same size were simulated since many existing methods cannot handle networks of different sizes.

### Study 1: Random network model with ground truth

Initial data vector *b_i_* at node *i* was simulated as independent and identically distributed multivariate normal across *n* subjects, i.e., 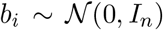 with the identity matrix *I_n_* as the covariance matrix of size *n × n*. Then, the new data vector *x_i_* at node *i* was generated by introducing additional dependency structures to *b_i_* through a mixed-effects model that partitions the covariance matrix of *x_i_* into *c* blocks forming modular structures [75] (Figure 7):

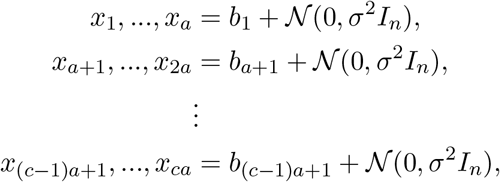

where *a* is the number of nodes in each module. In our simulation, we used *c* = 2, 5, 10, 20 and *a* = 50, 20, 10, 5 respectively, which give exactly *ca* = 100 nodes across networks. *σ* = 0.1 was universally used as network variability. Then, we computed Pearson correlation coefficient 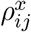 between *x_i_* and *x_j_*, which was then translated and scaled as 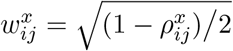 as a metric [24]. This gives a block network 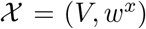. The mixed-effects model allows us to explicitly simulate the amount of statistical dependency between modules and nodes, providing the control over topological structures of connectedness.

**Figure 7:**
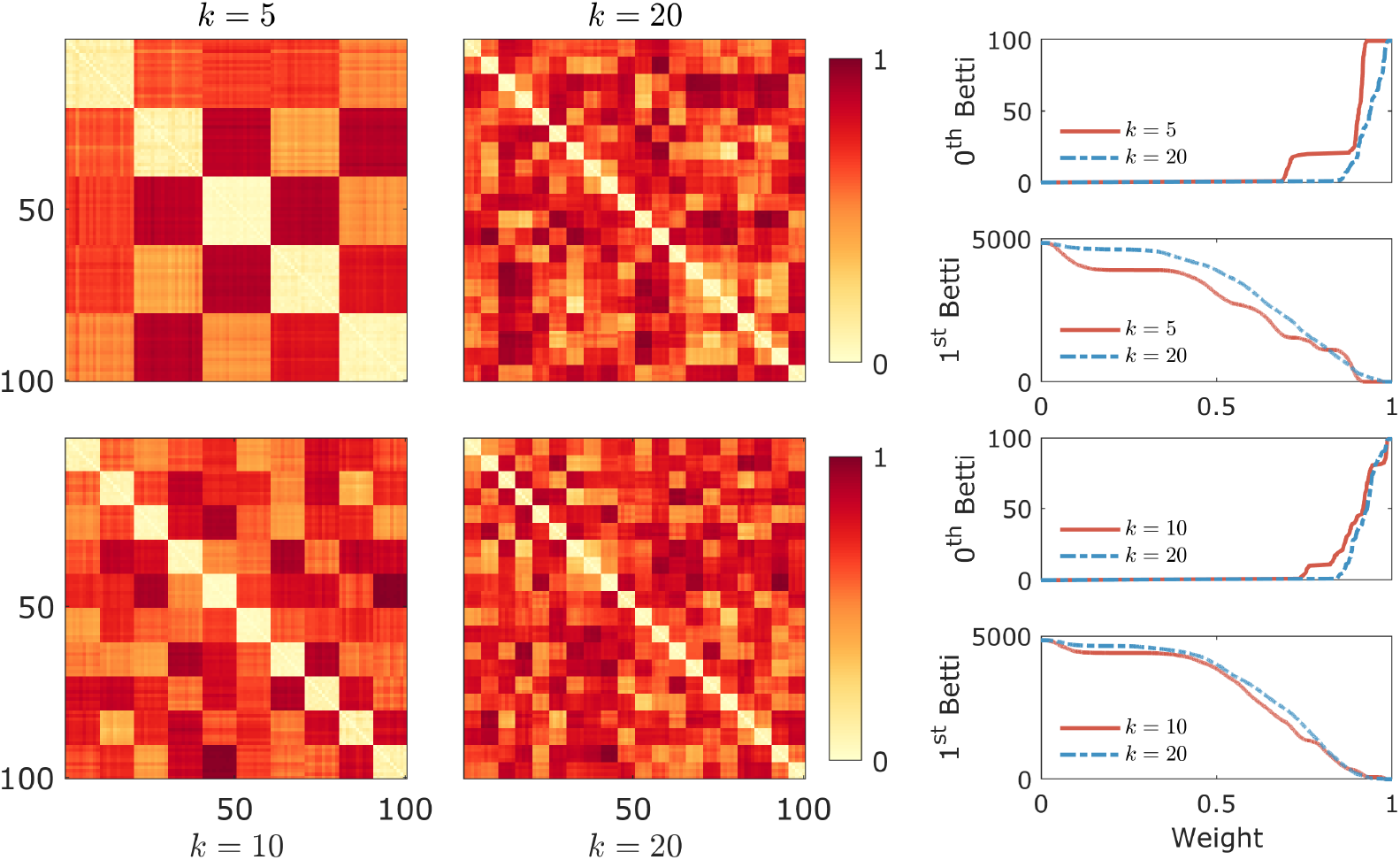
Study 1. Network difference. The comparison of networks with different topology: *k* = 5 vs. 20 (first row) and 10 vs. 20 (second row). The change of Betti numbers over filtration values clearly shows topology difference. The topological difference in the second row is subtle compared to the first row.

Based on the statistical model above, we simulated two groups of networks consisting of *n* = 7 subjects in each group for two different studies. We then tested the performance of topological loss for network differences. For comparison, we tested the topological loss against widely used Euclidean losses such as 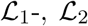 and 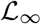-norms. We also compared against other topological distances such as bottleneck, Gromov-Hausdorff (GH) and Kolmogorov-Smirnov (KS) distances [76, 64, 77]. The bottleneck and GH-distance are two widely-used baseline distances in persistent homology often used in persistent diagrams and dendrograms [78] and brain networks [44]. KS-distance based on *β*_0_ and *β*_1_ curves is later introduced as a more intuitive alternative that gives results that are easier to interpret [79]. KS-distance was successfully applied to perform statistical inference on both *β*_0_ and *β*_1_ in a large-scale brain network studies [24]. For KS-distance, we computed its probability distribution exactly [34]. For all other distance, the permutation test was used. Since the sample size is small, the exact permutation test can be done by generating exactly 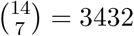 number of every possible permutation by shuffling the group labels to determine the empirical distribution of the two-sample *t*-statistic under the null.

#### Network difference

We compared networks generated by varying parameter *c* = 2 vs. 10, 5 vs. 20 and 10 vs. 20. Since the networks had different topological structures, the distances were expected to detect the differences (Figure 7). The simulations were independently performed 50 times and the performance results were given in terms of the false negative rate computed as the fraction of 50 simulations that gave *p*-values above 0.05 (Table 1). As expected, topological loss performed very well in all settings while other distances were less sensitive to more subtle topological difference. Even though the KS-distance on cycles (*β*_1_) performed better, it is known to be overly sensitive distance [24] and often produces false positives as shown in the next simulation.

**Table 1:** Study 1. The performance results are summarized in terms of false negative rate (top rows) and false positive rate (bottom rows).

| $c$ | $\mathcal{L}_1$ | $\mathcal{L}_2$ | $\mathcal{L}_\infty$ | GH | Bottleneck | KS($\beta_0$) | KS($\beta_1$) | $\mathcal{L}_{top}$ |
| --- | --- | --- | --- | --- | --- | --- | --- | --- |
| 2 vs. 10 | 0.02 | 0.00 | 0.02 | 0.30 | 0.40 | 0.00 | 0.00 | 0.02 |
| 5 vs. 20 | 0.06 | 0.00 | 0.02 | 0.20 | 0.32 | 0.02 | 0.00 | 0.00 |
| 10 vs. 20 | 1.00 | 0.86 | 0.34 | 0.32 | 0.20 | 0.24 | 0.00 | 0.08 |
| 2 vs. 2 | 0.00 | 0.00 | 0.00 | 0.00 | 0.00 | 0.54 | 0.88 | 0.00 |
| 5 vs. 5 | 0.00 | 0.00 | 0.00 | 0.00 | 0.00 | 0.10 | 0.42 | 0.00 |
| 10 vs. 10 | 0.00 | 0.00 | 0.00 | 0.00 | 0.00 | 0.02 | 0.12 | 0.00 |

#### No network difference

We compared networks generated by unvarying parameter *c* = 2 vs. 2, 5 vs. 5 and 10 vs. 10, which should give networks of similar topology in each group. It was expected that the networks were not topologically different and we should not detect the network differences. The simulations were independently performed 50 times and the performance results were given in terms of the false positive rate computed as the fraction of 50 simulations that gave *p*-values below 0.05 (Table 1). Except KS-distance, all the distances performed well.

While all the loss functions performed well when there was no network differences, the Euclidean losses, GH and bottleneck distances had the tendency to produce false negatives when there was network difference. On the other hand, KS-distance tended to produce false positives when there is no network difference. Overall, the proposed topological loss performed well.

### Study 2: Comparison against graph matching

The aim of this simulation is to evaluate the performance of the proposed topological matching process against existing graph matching algorithms [80, 81, 82, 83, 84] in differentiating networks of different sizes and topology. Graph matching algorithms are considered as the *baseline* for establishing correspondence between graphs and are usually done by penalizing network structures that cannot be exactly matched. Let *G*_1_ = (*V*_1_, *w*^1^) and *G*_2_ = (*V*_2_, *w*^2^) be two networks. For a graph matching problem modified for our brain network setting, where the edge weights are not binary but weighted, we need to find mapping *τ_gm_* between nodes *i*_1_, *j*_1_ ∈ *V*_1_ and *i*_2_, *j*_2_ ∈ *V*_2_ that best preserves edge attributes between edge weights 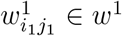 and 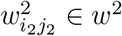 words, we seek *τ_gm_* that maximizes the graph matching cost

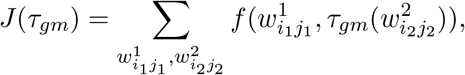

where *f* measures the similarity between edge attributes and the summation is taken over all possible edge weights. The matching cost *J* (*τ_gm_*) quantifies similarity between two networks by taking large values for similar networks and value close to zero for dissimilar networks hence *J* (*τ_gm_*) is somewhat the inverse of distance metrics. We compared the proposed topological loss against four wellknown graph matching algorithms: graduated assignment (GA) [80], spectral matching (SM) [81], integer projected fixed point method (IPFP) [83] and reweighted random walk matching (RRWM) [82]. Such graph matching methods are widely used as baseline algorithms in medical imaging, computer vision and machine learning studies [84, 85, 86, 87, 88, 89]. For all the baseline methods, we used existing implementation codes from authors’ repository websites listed in the publication. We also used parameters recommended in the public code for each baseline algorithm *without* modification. Since we are dealing with weighted edges, the graph matching algorithms based on binary edge weight such as [68] are excluded in the study.

In study 2, a different random network model from study 1 is used. We simulate random modular network 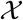 with *d* number of nodes and *c* number of modules, where the nodes are evenly distributed among modules. Figure 8 displays large modular networks with *d* = 120 nodes and *c* = 2, 3, 6 modules such that we have *d/c* = 60, 40, 20 number of nodes in each module, respectively. Since time complexity of the aforementioned graph matching algorithms can be very demanding (Figure 9), we considered *d* = 12, 18, 24 and *c* = 2, 3, 6 in this simulation. Each edge connecting two nodes within the same module was then assigned a random weight following a normal distribution 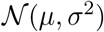 with probability *p* or otherwise Gaussian noise 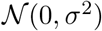 with probability 1*− p*. On the other hand, edge weights connecting nodes between different modules had probability 1 −*p* of being 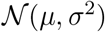 and probability *p* of being 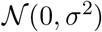. With larger value of within-module probability *p*, we have more pronounced modular structure. Any negative edge weights were set to zero. This gives the random network 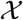 that exhibits topological structures of connectedness. Figure 8 illustrates changes of network modular structure as parameters *p* and *c* vary. We used *µ* = 1 and *σ* = 0.25 universally throughout study 2.

**Figure 8:**
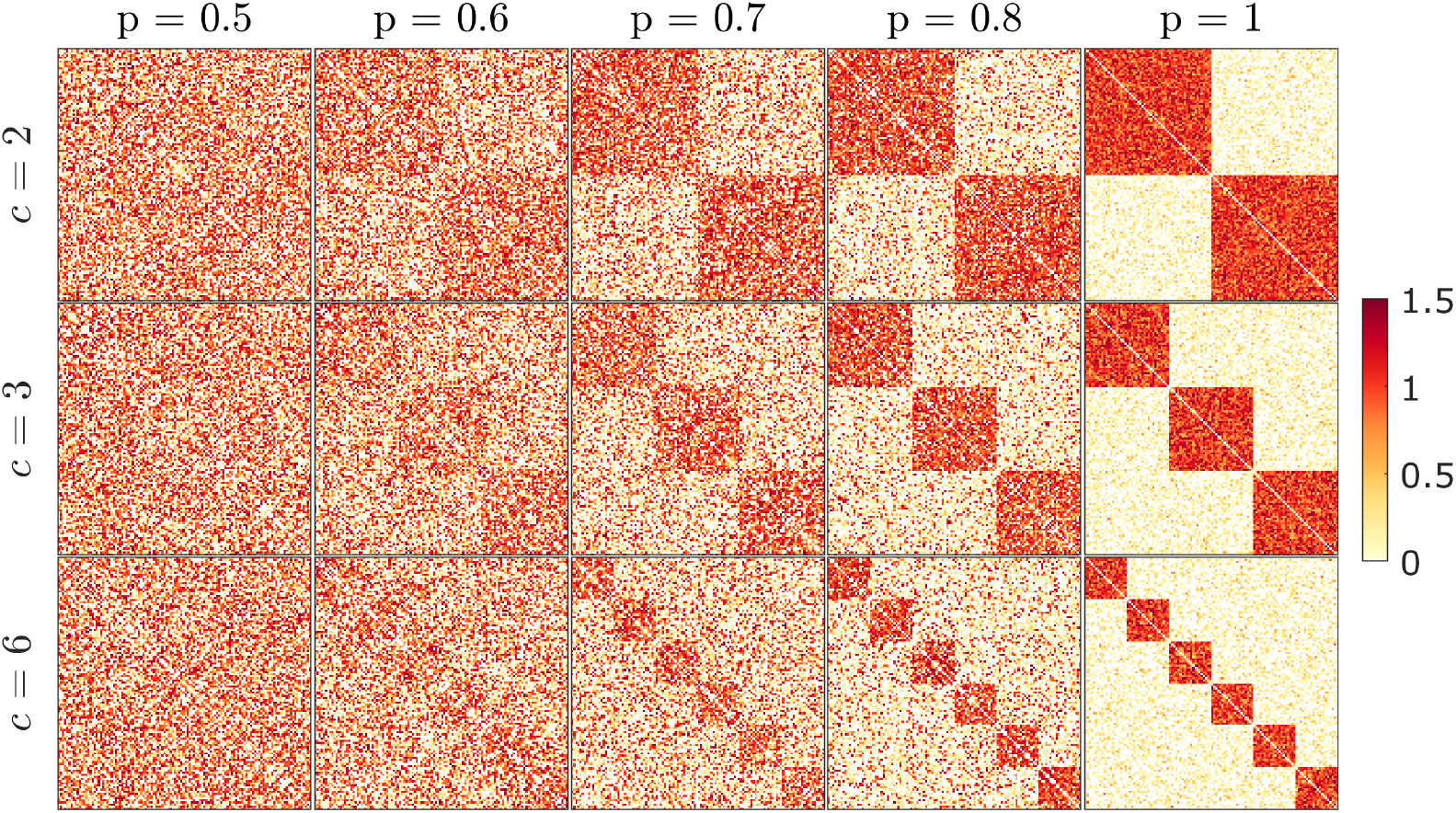
Study 2 simulation examples. Network modular structure varies as parameters *p* (probability of connection within modules) and *c* (the number of modules) change. The modular structure becomes more pronounced as *p* increases.

**Figure 9:**
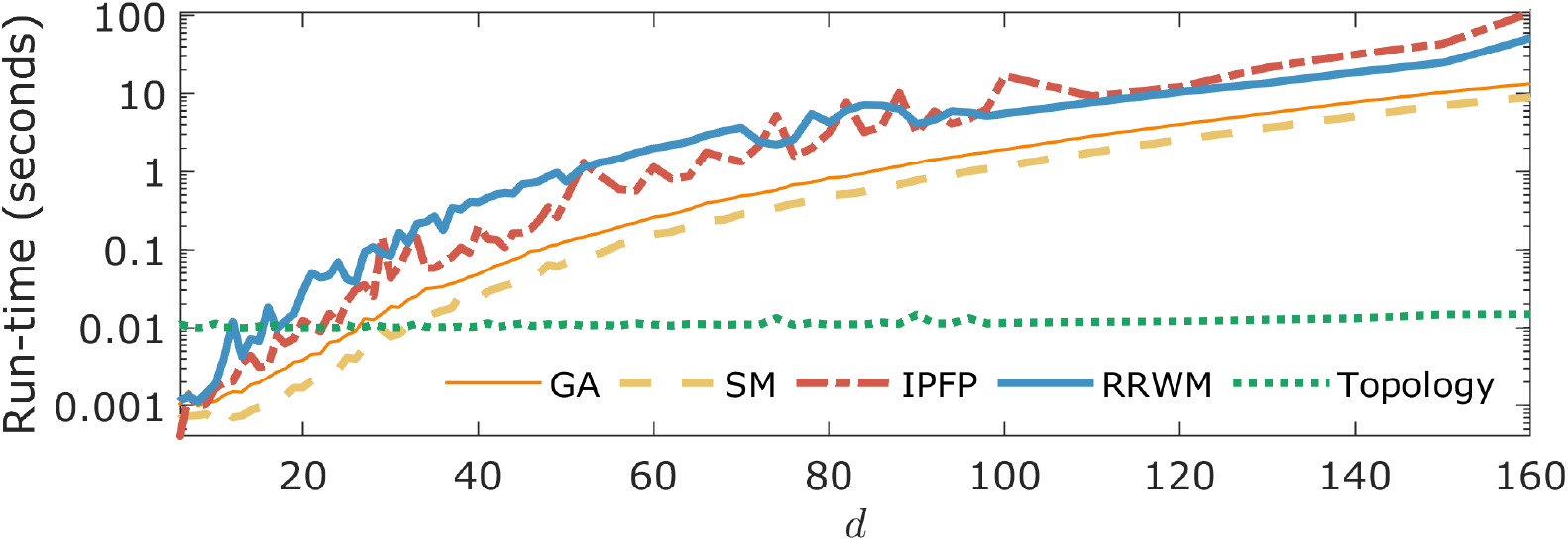
Study 2 run-time. The amount of time each algorithm takes to compute its matching cost between two modular networks of size *d*. Average run-times of topological loss and other baseline graph matching algorithms are plotted over *d*. The run-time performance of the baseline methods is consistent with [85] for GA and SM, and [88] for IPFP and RRWM.

Based on the statistical model above, we simulated two groups of random modular networks 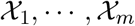 and 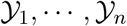. If there is group difference, the topological loss is expected to be relatively small within groups and relatively large between groups. The average topological loss within group given by

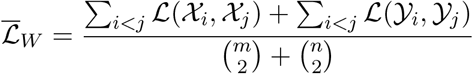

is expected to be smaller than the average topological loss between groups given by

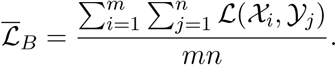

We measure the disparity between groups as the ratio statistic 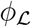

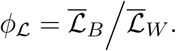

If 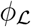 is large, the groups differ significantly in network topology. On the other hand, if 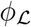 is small, it is likely that there is no group differences. Similarly, we define the ratio statistic for graph matching cost *J* as

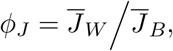

where 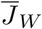 is the average graph matching cost within groups and 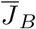 is the average graph matching cost between groups. Since the distributions of the ratio statistics 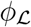 and *φ_J_* are unknown, the permutation test is used to determine the empirical distributions. Figure 10 displays the empirical distribution of 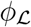. By comparing the observed group ratio 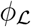 to this empirical distribution, we can determine the statistical significance of testing the group difference. However, when the sample size is large, existing matching algorithms are slow for the permutation test. So we adapted for a scalable computation strategy as follows [18].

**Figure 10:**
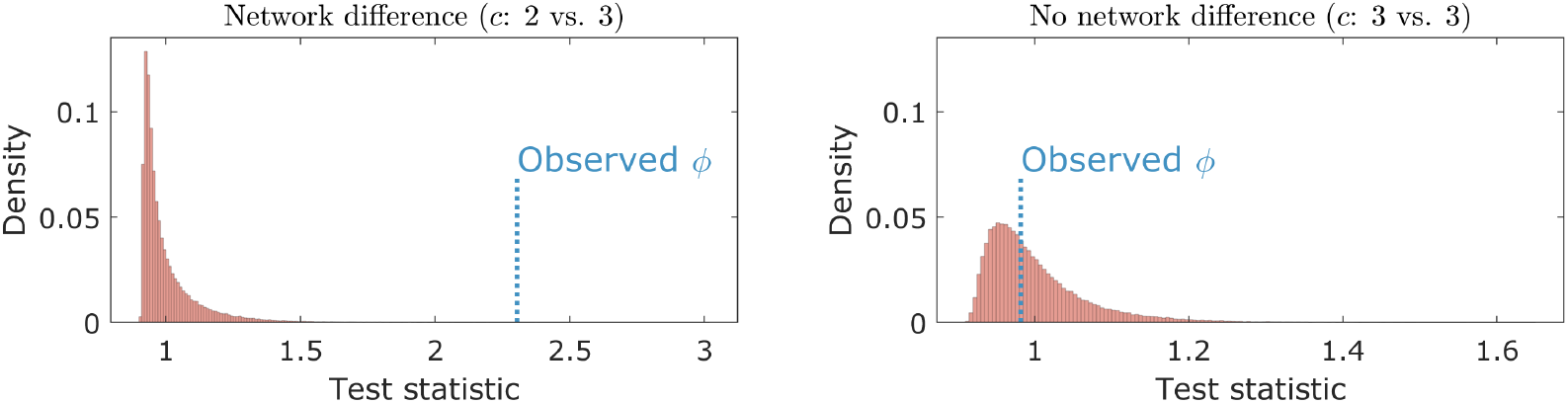
Empirical distributions in Study 2. The empirical distributions of the ratio statistic were generated by the permutation on two groups each consisting of 10 modular networks. Here we tested if there is group difference in networks with varying parameter *c* = 2 vs. 3 (left) and 3 vs. 3 (right). As expected, the test based on the topological loss rejected the null hypothesis when there is group difference.

Given two groups of networks, topological loss (or graph matching cost) for every pair of networks needs to be computed only once, which can then be arranged into a matrix whose rows and columns are networks, and the *ij*-entry is the loss between two networks corresponding to row *i* and column *j* (Figure 11). Once we obtain such matrix, the permutation process is equivalent to rearranging rows and columns based on permuted group labels. There are 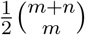 total number of permutations excluding the symmetry of loss functions. Computing the ratio statistic over a permutation requires re-summing over all such losses, which is time consuming. Instead, we can perform the transposition procedure of swapping only one network per group and setting up iteration of how the ratio statistic changes over the transposition [18].

**Figure 11:**
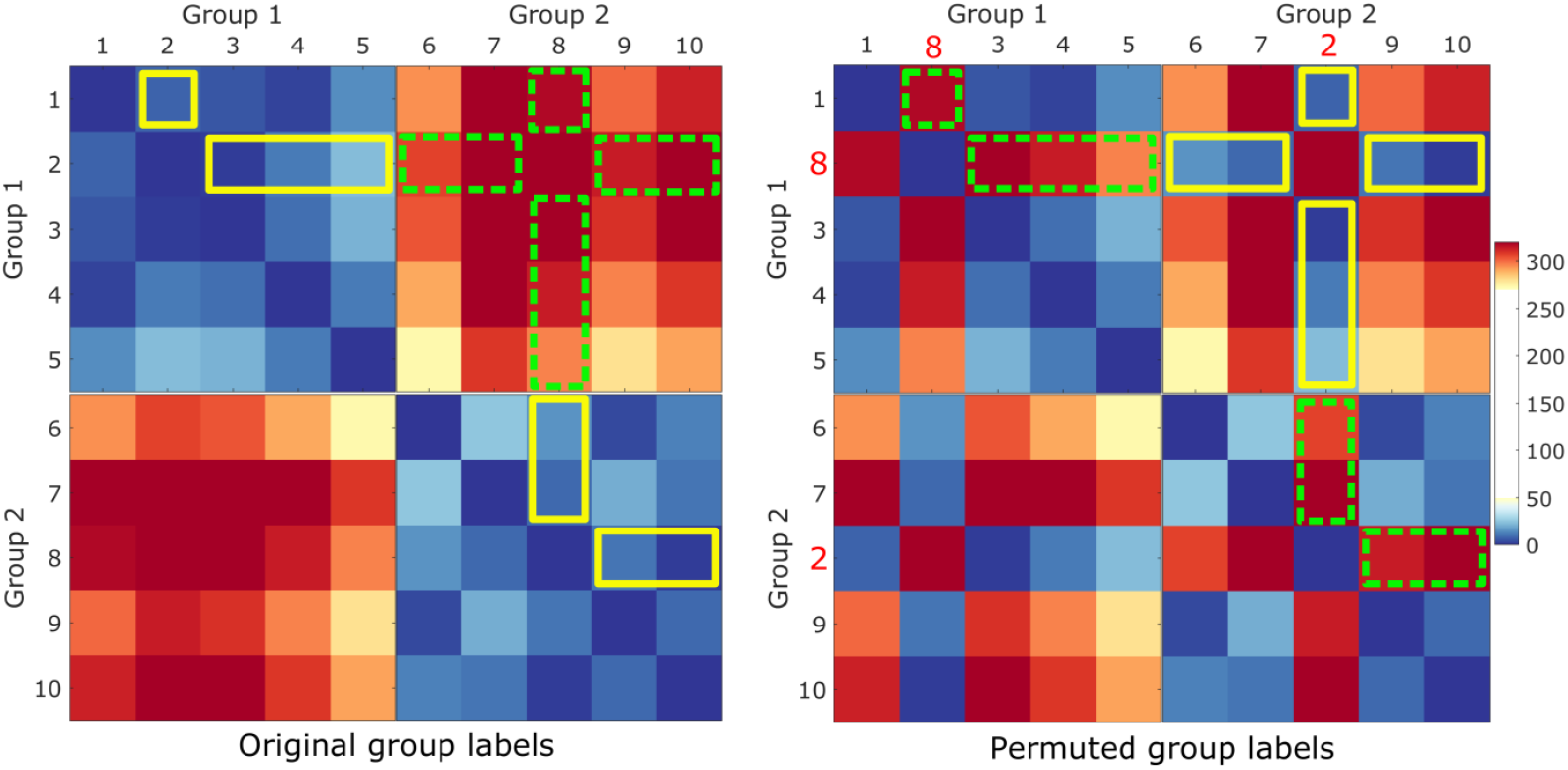
Study 2. Two groups each consisting of 5 modular networks simulated with parameters *c* = 2 vs. 3, resulting in small and large topological losses within groups and between groups respectively. Left: the matrix whose *ij*-th entry represents the loss between network *i* and *j*. The main diagonal consists of zero since topological loss between two identical networks is zero. Right: a transposition between the 2-nd network in Group 1 and the 8-th network in Group 2. We do not need to recompute all the pairwise losses again but just rearrange them. In particular, only the pairwise losses in the yellow solid lines and green dashed lines are rearranged. Thus, we simply need to figure out how the rearrangement of entries changes the ratio statistic in an iterative manner. This enables us to easily perform the permutation test in a scalable fashion. We compute *δ* in equation (4) by subtracting the sum of entries within the yellow lines from the sum of entries within the green lines.

Let 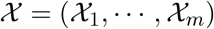 and 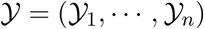 be the two groups of networks. We *transpose k*-th and *l*-th networks between the groups as

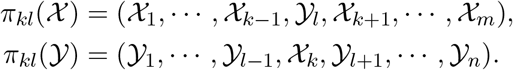

Over transposition *π_kl_*, the ratio statistic is changed from 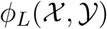 to 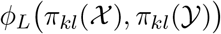 which involves the following functions:

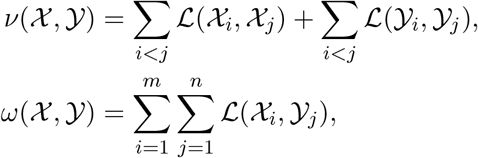

where *ν* is the total sum of within-group losses and *ω* is the total sum of between-group losses. Then, we determine how *ν* and *ω* change over the transposition *π_kl_*. As 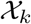 and 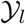 are swapped, the function *ν* is updated over the transposition *π_kl_* as (Figure 11)

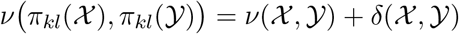

with

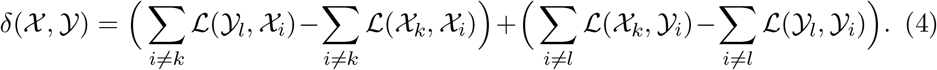

Similarly, function *ω* are updated iteratively over the transposition *π_kl_* as:

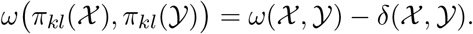

The ratio statistic over the transposition is then computed as

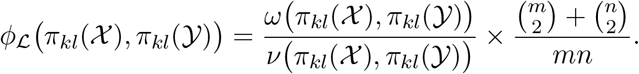

For each transposition, we store information about the function values *ν* and *ω*, and update them sequentially. Each transposition requires manipulating 2(*m* + *n* − 2) terms as opposed to 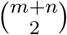 total number of terms over a random permutation. More transpositions than permutations can be generated given the same amount of run-time, which speed up the convergence of transposition procedure [18]. To further accelerate the rate of convergence and avoid possible bias, we introduce a full permutation to the sequence of transpositions. Figure 12 illustrates the convergence of transposition procedure.

**Figure 12:**
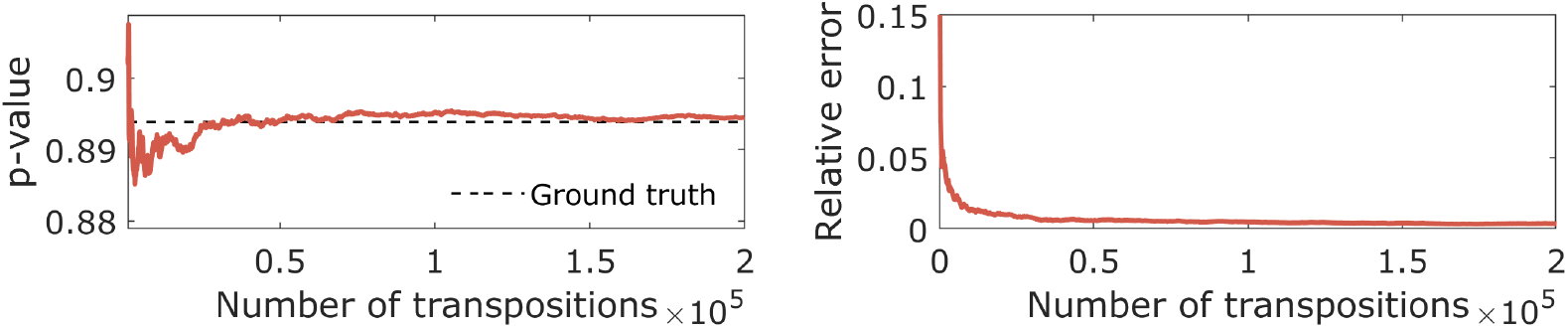
Transposition test is applied in determining the statistical significance of two groups each consisting of 10 simulated networks. To further speed up the convergence rate, a random permutation is intermixed for every sequence of 500 transpositions. The left panel shows the convergence of *p*-value over 50000 transpositions. For comparison, ground truth *p*-value is computed from the exact permutation test by enumerating every possible permutations exactly once. The left panel shows the average relative error against the ground truth across 100 independent simulations.

In each simulation, we generated two groups each with 10 random modular networks. We then computed 200000 sequentially random transpositions while interjecting a random permutation for every 500 transpositions and obtained the *p*-values. This guarantees the convergence of *p*-values within 2 decimal places (within 0.01) in average. The simulations were independently performed 50 times and the average *p*-value across 50 simulations was reported.

#### Network difference

We compared two groups of networks generated by varying parameter *c* = 2 vs. 3, 2 vs. 6 and 3 vs. 6 each with *d* = 12, 18, 24 nodes and *p* = 0.6, 0.8 probability of connection within modules. Since each group exhibited different modular network structure, topological loss and graph matching cost were expected to detect the group difference. Table 2 summarizes the performance results. Networks with *p* = 12 nodes might be too small to extract topologically distinct features used in each algorithm. Thus, all graph matching costs performed poorly while the topological loss performed reasonably well. When the number of nodes increases, all methods show overall performance improvement. In all settings, topological loss significantly outperforms other graph matching algorithms.

**Table 2:**
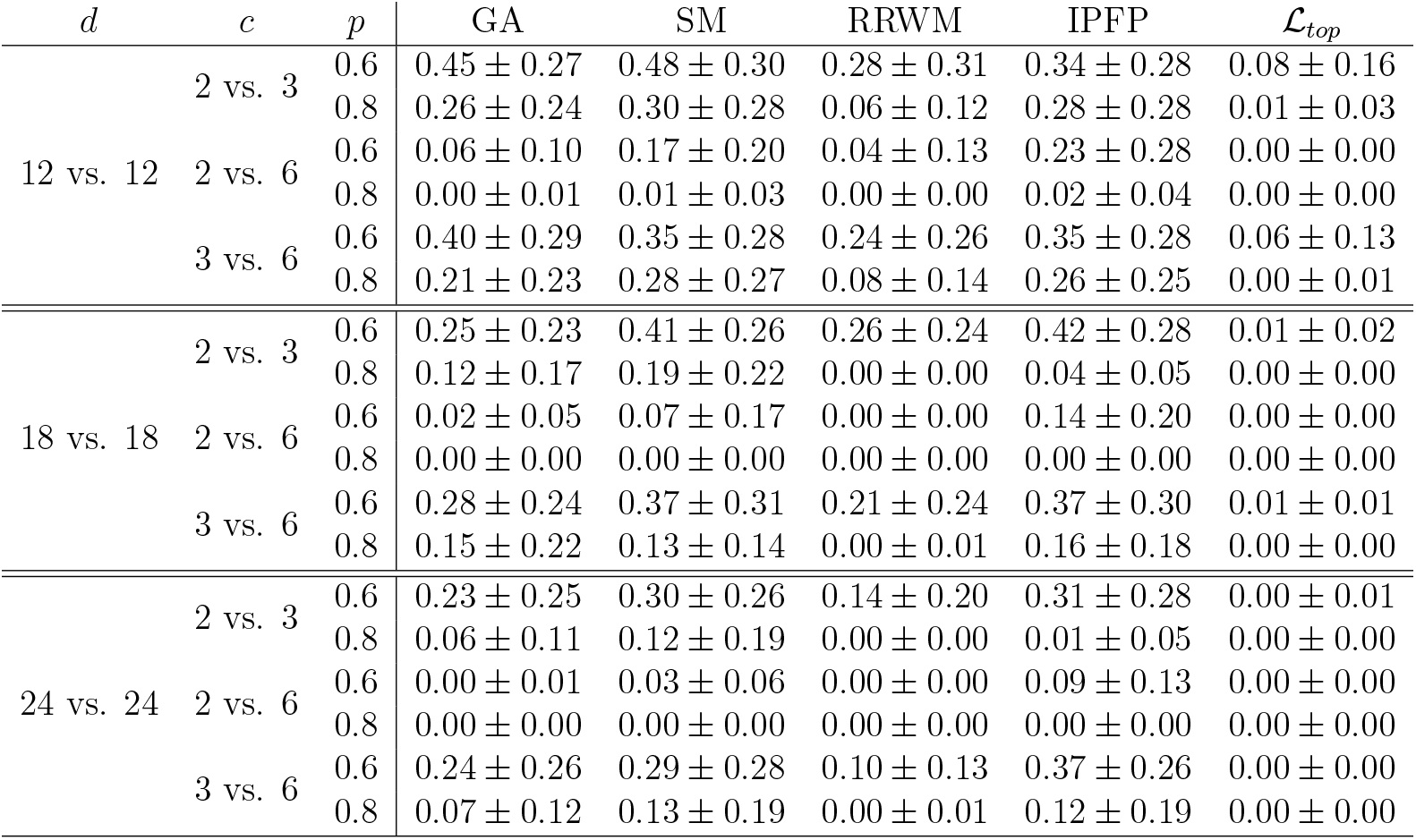
Study 2. Network difference. The performance results are summarized as average *p*-values for various parameter settings of *d* (number of nodes), *c* (number of modules) and *p* (probability of connection within modules).

#### No network difference

We compared networks generated by unvarying parameter *c* = 2 vs. 2, 3 vs. 3 and 6 vs. 6 each with *d* = 12, 18, 24 nodes and *p* = 0.6, 0.8 probability of connection within modules. Since it was expected that there was no topological difference between networks generated using the same values for parameters *c, d* and *p*, topological loss and graph matching cost should not be able to detect the group difference. The performance result is summarized in Table 3. In all settings, all methods performed well when there was no group difference.

**Table 3:**
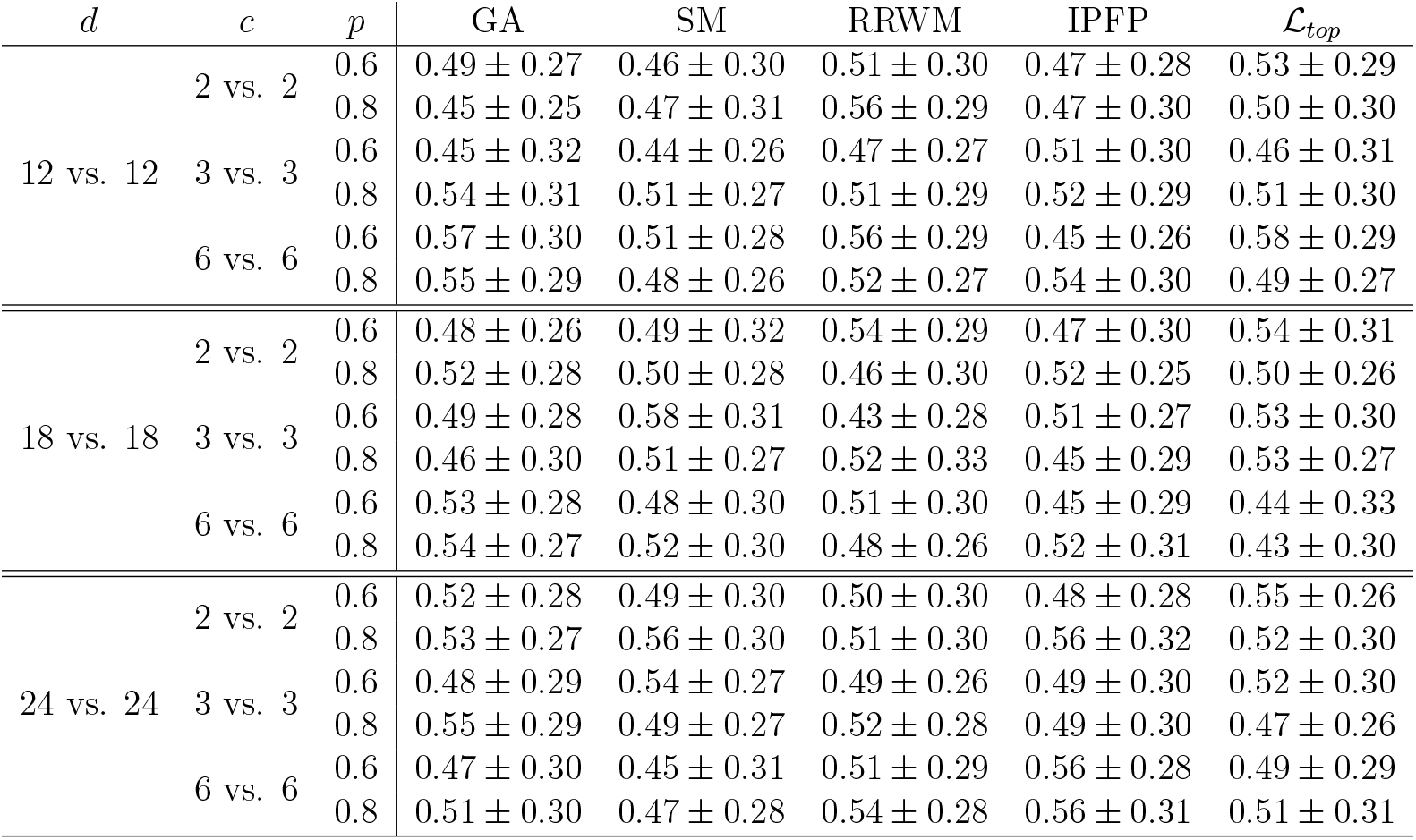
Study 2. No network difference. The performance results are summarized as average *p*-values for various parameter settings of *d* (number of nodes), *c* (number of modules) and *p* (probability of connection within modules).

The baseline graph matching methods have low sensitivity over topological differences such as connected components and cycles in networks. They were unable to detect the network differences, especially subtle topological difference. While it might be possible to extend a graph matching algorithm to encode high-order geometrical relations, a small increment in the order of relations usually result in a combinatorial explosion of the amount of data needed to fit the model [84]. Thus, most of high-order graph matching methods is often limited to very sparse networks such as binary trees. They are not practical in dense functional brain networks with far larger number of cycles. On the other hand, the proposed topological loss is able to detect such subtle topological pattern differences with minimal amount of run-time.

## 4 Application

### Dataset and preprocessing

The dataset is the resting-state fMRI (rsfMRI) of 412 subjects collected as part of the Human Connectome Project (HCP) twin study [52, 53]. The rsfMRI were collected over 14 minutes and 33 seconds using a gradient-echo-planar imaging (EPI) sequence with multiband factor 8, repetition time (TR) 720 ms, time echo (TE) 33.1 ms, flip angle 52^*◦*^, 104 90 (RO PE) matrix size, 72 slices, 2 mm isotropic voxels, and 1200 time points. Subjects without rsfMRI or full 1200 time points were excluded. Additional details on the imaging protocol is given in https://protocols.humanconnectome.org/HCP/3T/imaging-protocols.html. During each scanning, participants were at rest with eyes open with relaxed fixation on a projected bright cross-hair on a dark background [53]. The standard minimal preprocessing pipelines [90] were applied on the fMRI scans including spatial distortion removal [91, 92], motion correction [93, 94], bias field reduction [95], registration to the structural MNI template, and data masking using the brain mask obtained from FreeSurfer [90]. This resulted in the resting-state functional time series with 91 *×* 109 *×* 91 2-mm isotropic voxels at 1200 time points. The subject ranged from 22 to 36 years in age with average age 29.24 *±* 3.39 years. There are 172 males and 240 females. Among them, there are 131 MZ-twin pairs and 75 same sex DZ-twin pairs.

Subsequently, we employed the Automated Anatomical Labeling (AAL) template to parcellate the brain volume into 116 non-overlapping anatomical regions [2]. We averaged fMRI across voxels within each brain parcellation, resulting in 116 average fMRI time series with 1200 time points for each subject. Previous studies reported that head movement produces spatial artifacts in functional connectivity [96, 97, 98, 99]. Thus, we scrubbed the data to remove fMRI volumes with significant head motion [96, 100]. We calculated the framewise displacement (FD) from the three translational displacements and three rotational displacements at each time point to measure the head movement from one volume to the next. The volumes with FD larger than 0.5 mm and their neighbors (one back and two forward time points) were scrubbed [97, 96, 100]. We excluded 12 subjects having excessive head movement, resulting in fMRI data of 400 subjects (168 males and 232 females). More than one third of 1200 volumes being scrubbed in the excluded 12 subjects. Among the remaining 400 subjects, there are 124 monozygotic (MZ) twin pairs and 70 same-sex dizygotic (DZ) twin pairs. The first 20 time points were removed from all subjects to avoid artifacts in the fMRI data, leaving 1180 time points per subject [101, 102].

In [18], the white matter fiber orientation information was extracted by multishell, multi-tissue constrained spherical deconvolution from different tissue types such as white matter and gray matter [103, 104]. The fiber orientation distribution functions were estimated and apparent fiber densities were exploited to produce the reliable white and gray matter volume maps [104, 105]. Subsequently, multiple random seeds were selected in each voxel to generate about 10 million initial streamlines per subject with the maximum fiber tract length at 250 mm and FA larger than 0.06 using MRtrix3 (http://www.mrtrix.org) [106, 107]. The Spherical-Deconvolution Informed Filtering of Tractograms (SIFT2) technique making use of complete streamlines was subsequently applied to generate more biologically accurate brain connectivity, which results in about 1 million tracts per subject [108]. Nonlinear diffeomorphic registration between subject images to the template was performed using ANTS [109, 110]. AAL was used to parcellate the brain into 116 regions [2]. The subject-level connectivity matrices were constructed by counting the number of tracts connecting between brain regions. The structural brain network *P* that serves as the template where all the functional networks are aligned is obtained by computing the one sample *t*-statistic map *P* over all the subjects and rescaling *t*-statistics between 0 to 2 using the hyperbolic tangent function tanh, then adding 1.

### Learning individual networks

Among 400 subjects, there are *p* = 124 monozygotic (MZ) twin pairs and *q* = 70 same-sex dizygotic (DZ) twin pairs. For subject *k*, we have resting-state fMRI time series *x* = (*x*_1_, *x*_2_, *…, x*_1180_) for region *i* and *y* = (*y*_1_, *y*_2_, *…, y*_1180_) for region *j* with 1180 time points. Correlation 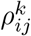 between regions *i* and *j* was computed as the Pearson correlation between *x* and *y*. This gives the correlation matrix 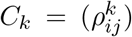 which is used as the baseline against the proposed method. We then translate and scale the correlation 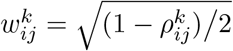, which is a metric [24]. The subject-level functional brain network is given by *G_k_* = (*V, w^k^*)The *t*-statistic map *P* is used as the template structural brain network where the functional network *G_k_* is matched.

Given *λ_k_*, we applied the topological learning to estimate the subject-level network Θ_*k*_(*λ_k_*) by minimizing the objective (1) using the individual network *G_k_* and the structural network *P*:

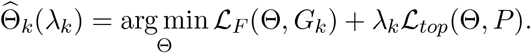

For every subject, we globally used *λ* = 1.0000 where the average minimum is obtained (Figure 3).

### Heritability in twins

We investigated in localizing what part of brain networks are genetically heritable. In particular, we investigated if the estimated network 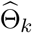 is genetically heritable in twins as follows. At edge *ij*, let (*a*_1*l*_, *a*_2*l*_) be the *l*-th twin pair in MZ-twin and (*b*_1*l*_, *b*_2*l*_) be the *l*-th twin pair in DZ-twin. MZ-twin and DZ-twin pairs are then represented as

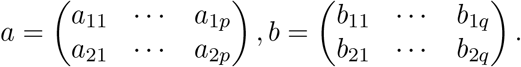

Let *a_r_* = (*a_r_*_1_, *a_r_*_2_, *…, a_rp_*) and *b_r_* = (*b_r_*_1_, *b_r_*_2_, *…, b_rq_*). Then, MZ-correlation is computed as the Pearson correlation *γ^MZ^* (*a*_1_, *a*_2_) between *a*_1_ and *a*_2_. Similarly for DZ-correlation *γ^DZ^* (*b*_1_, *b*_2_). In the widely used ACE genetic model, the heritability index (HI) *h*, which determines the amount of variation caused by genetic factors in population, is estimated using Falconer’s formula [111, 24]. Thus, HI *h* is given by *h*(*a, b*) = 2(*γ^MZ^ − γ^DZ^*).

Since the order of the twins is interchangeable, we can *transpose* the *l*-th twin pair in MZ-twin as

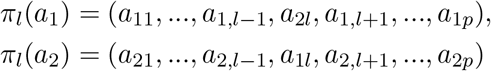

and obtain another MZ-correlation *γ^MZ^* (*π_l_*(*a*_1_), *π_l_*(*a*_2_)). Likewise, we obtain many different correlations for DZ-twin. Similar to the transposition test used in the simulation study 2, we perform a sequence of random transpositions iteratively to estimate the twin correlations *γ^MZ^* and *γ^DZ^* sequentially as follows [18].

Over the transposition *π_l_*, the MZ-correlation is changed from *γ^MZ^*(*a*_1_, *a*_2_) to *γ^MZ^* (*π_l_*(*a*_1_), *π_l_*(*a*_2_)), which involves the following functions:

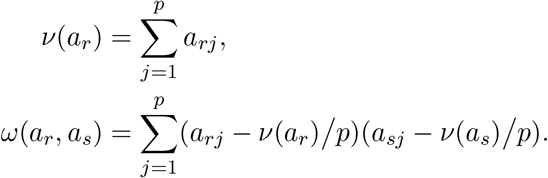

The functions *ν* and *ω* are updated iteratively over the transposition *π_l_* as:

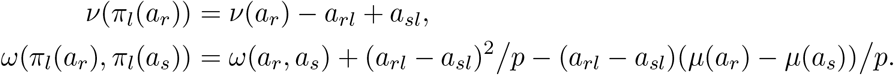

Then, the MZ-correlation after transposition is calculated by

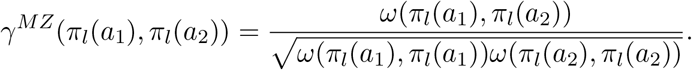

The time complexity for computing correlation iteratively is 33 operations per transposition, which is significantly more efficient than that of direct correlation computation per permutation. In numerical implementation, we sequentially perform random transpositions *π_l_*1, *π_l_*2, *…, π_l_J*, which result in *J* different twin correlations. Let

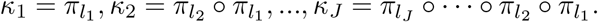

The average MZ-correlation 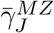 of *J* correlations is given by

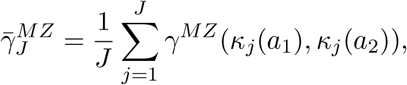

which is iteratively updated as:

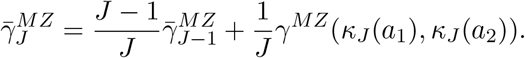

The average correlation 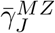 converges to the true underlying twin correlation *γ^MZ^* for sufficiently large *J*. Similarly, DZ-correlation *γ^DZ^* is estimated.

### Results

Using the transposition method, we randomly transposed a twin and updated the correlation for 50000 times. This process was repeated 100 times and the total 50000 *×* 100 correlations were used in estimating the underlying MZ- and DZ-correlations. At every edge, the standard deviation of the average correlations from 100 results was smaller than 0.01, which guarantees the convergence of the estimate within two decimal places in average.

We computed the HI-maps using the original correlation matrix *C_k_* and the proposed topologically learned network 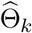. If such method is more sensitive toward identifying network phenotypes, it is expected that we detect more high HI values along edges. Figure 13 shows far more connections with 100% heritability for the topologically learned network 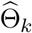 compared to the original Pearson correlation matrix *C_k_*. The network 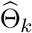 is expected to inherit topological sparsity from the template structural brain network *P* that has sparse topology with less cycles (Figure 5). This suggests that noisy, short-lived cycles were removed from the functional networks, improving the statistical sensitivity. Comparing HI-maps from the two methods, there are overlaps but the topologically learned approach detects more connections with higher heritability. For network 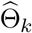, left superior parietal lobule and left amygdala connection shows the strongest heritability among many other connections (Table 4).

**Figure 13:**
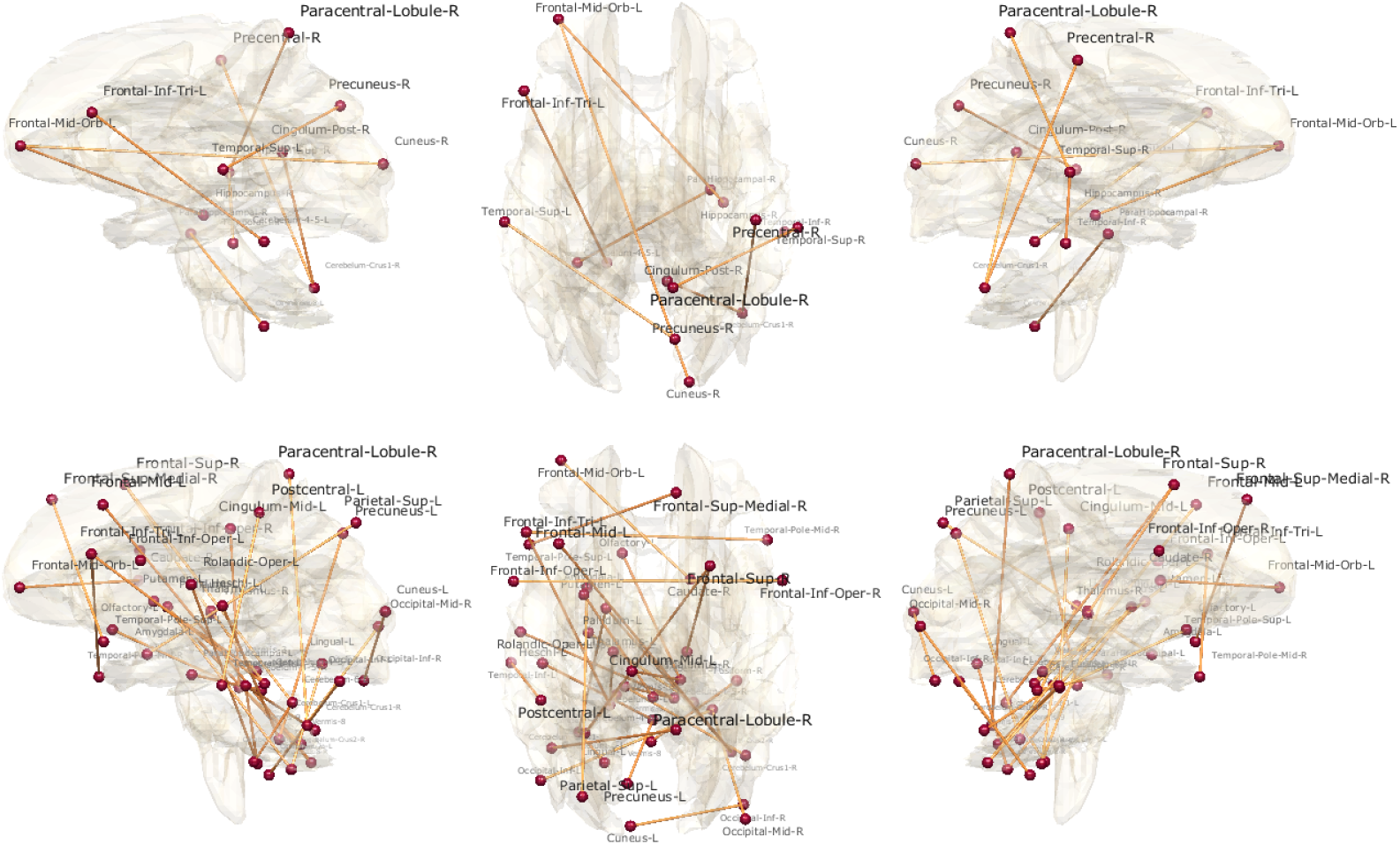
Most highly heritable connections. The heritability index maps display only connections with 100% heritability (HI ≥1) using the original Pearson correlation matrix (top) and the topologically learned network (bottom).

**Table 4:**
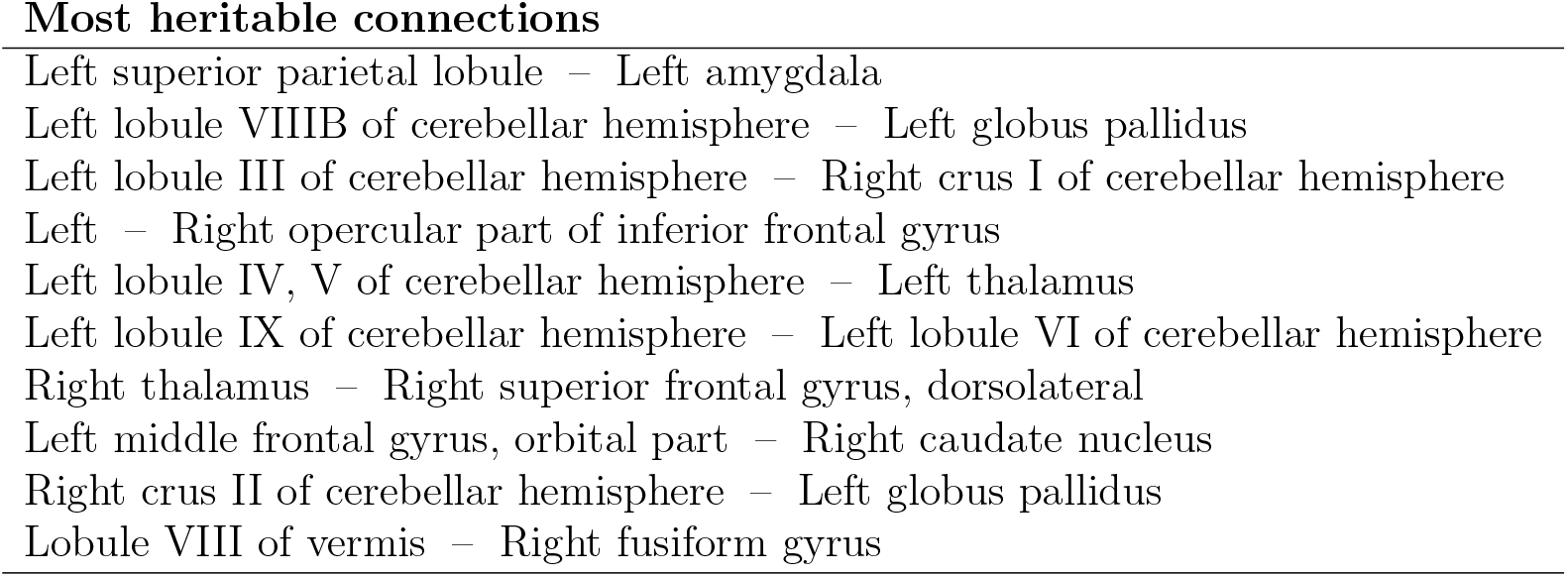
Top ten most heritable connections with 100% heritability (HI ≥ 1) using the topologically learned network. Such connections are the top ten highest HI’s among other connections with 100% heritability.

## 5 Conclusion

We have presented a novel topological learning framework that can analyze networks of different sizes and topology through a new loss function. Among many different learning tasks, the method is illustrated with averaging and regression. The method is briefly shown on how to average networks of different sizes and topology, which is not easy with existing methods. We have also applied the method to twin brain imaging data in analyzing functional and structural brain networks together.

Our topological learning framework is more sensitive on detecting subtle genetic network signals than the baseline method. The topological loss can be easily adapted to other learning tasks such as clustering. It is well known that the Euclidean-distance based clustering such as *k*-means clustering does not perform well against more geometric clustering method such as spectral clustering [112, 113]. It may be possible that a new clustering method utilizing the proposed loss function might perform better than *k*-means or spectral clustering. This is left as a future study.

## Acknowledgements

We thank Li Shen of University of Pennsylvania for providing the *t*-statistic map of structural brain networks used in this study. The *t*-statistic map came from study [114]. This study is funded by NIH R01 EB022856, EB02875, NSF MDS2010778.

